# DAF-16/FoxO and DAF-12/VDR control cellular plasticity both cell-autonomously and via interorgan signaling

**DOI:** 10.1101/2020.12.15.422925

**Authors:** Ulkar Aghayeva, Abhishek Bhattacharya, Surojit Sural, Eliza Jaeger, Matthew Churgin, Christopher Fang-Yen, Oliver Hobert

**Author notes:** equal contributions (Abhishek Bhattacharya, Surojit Sural).

## Abstract

Many cell types display the remarkable ability to alter their cellular phenotype in response to specific external or internal signals. Such phenotypic plasticity is apparent in the nematode *C. elegans* when adverse environmental conditions trigger entry into the dauer diapause stage. This entry is accompanied by structural, molecular and functional remodeling of a number of distinct tissue types of the animal, including its nervous system. The transcription factor effectors of three different hormonal signaling systems, the insulin-responsive DAF-16/FoxO transcription factor, the TGFβ-responsive DAF-3/SMAD transcription factor and the steroid nuclear hormone receptor, DAF-12/VDR, a homolog of the vitamin D receptor, were previously shown to be required for entering the dauer arrest stage, but their cellular and temporal focus of action for the underlying cellular remodeling processes remained incompletely understood. Through the generation of conditional alleles that allowed us to spatially and temporally control gene activity, we show here that all three transcription factors are not only required to initiate tissue remodeling upon entry into the dauer stage, as shown before, but are also continuously required to maintain the remodeled state. We show that DAF-3/SMAD is required in sensory neurons to promote and then maintain animal-wide tissue remodeling events. In contrast, DAF-16/FoxO or DAF-12/VDR act cell autonomously to control anatomical, molecular and behavioral remodeling events in specific cell types. Intriguingly, we also uncover non-cell autonomous function of DAF-16/FoxO and DAF-12/VDR in nervous system remodeling, indicating the presence of several insulin-dependent inter-organ signaling axes. Our findings provide novel perspectives on how hormonal systems control tissue remodeling.

## INTRODUCTION

The identity of a fully differentiated cell in a multicellular organism is usually described by a number of phenotypic criteria, ranging from overall anatomy to cellular function to molecular features. Once such differentiated state has been acquired, it is often thought to persist throughout the life of an animal and be controlled by active maintenance mechanisms (Blau, 1992). Nonetheless, Rudolf Virchow already pointed out in 1886 that there are “plastic processes” that accompany the transition of a differentiated cell into a different state, particularly in the context of disease (Mills et al., 2019). The plasticity of cellular identity has now become a widely accepted phenomenon, occurring in a number of different cellular and organismal contexts (Mills et al., 2019). In the brain, cellular plasticity phenomena become evident in a number of different contexts and include cellular remodeling events that occur after injury, after the encounter of stressful environmental conditions or during specific developmental transitions, such as puberty. Several hormonal signaling systems have been implicated in triggering a number of distinct structural and functional remodeling events in the vertebrate brain (McEwen, 2010).

Remarkable changes of cellular phenotypes within distinct tissue types, including the nervous system, are observed upon entry in the dauer stage, a diapause stage of rhabditid nematodes (Cassada and Russell, 1975). In response to detrimental environmental conditions perceived at a specific postembryonic larval stage, epidermal and muscle cell types shrink their volume, resulting in an overall constriction of animal shape (Androwski et al., 2017; Cassada and Russell, 1975; Golden and Riddle, 1982). The extracellular collagen cuticle, secreted by skin cells, is remodeled, and intestinal cells alter their metabolic state (Androwski et al., 2017; Burnell et al., 2005). Several profound cellular remodeling events are observed in the nervous system: Some sensory neurons grow extensive dendritic branches (Schroeder et al., 2013), some glial cells change their ensheathment of several neuron types (Albert and Riddle, 1983; Procko et al., 2011), and there is extensive remodeling of electrical synapses throughout the entire nervous system (Bhattacharya et al., 2019). The expression patterns of many G-protein coupled sensory receptors are altered in a highly specific manner in a number of sensory-, inter- and motor neuron types (Nolan et al., 2002; Peckol et al., 2001; Vidal et al., 2018). On an animal-wide level, there are global changes in gene expression and chromatin modification patterns (Hall et al., 2010), including wide-spread changes in neuropeptide gene expression (Lee et al., 2017). Paralleling these changes in cellular and molecular phenotypes, animals undergo a number of striking behavioral changes, from altered sensory responses to changes in locomotory and exploratory strategies to a complete silencing of the enteric system (Bhattacharya et al., 2019; Cassada and Russell, 1975; Gaglia and Kenyon, 2009; Lee et al., 2012). Most of these remodeling events are reversible. After the improvement of external conditions animals exit the dauer stage and develop into adults whose anatomy and behavior at least superficially resembles that of worms that have not passed through the dauer stage, even though some molecular changes appear to persist (Hall et al., 2010; Sims et al., 2016; Vidal et al., 2018).

Most cells that are remodeled upon entry into the dauer state have already been born and differentiated into fully functional cell types during embryogenesis and early larval development. The ability of postmitotic, differentiated cells to remodel a number of their anatomical, molecular and functional features during dauer entry in mid-larval development (and their ensuing reversibility upon dauer exit) is a remarkable example of the plasticity of the cellular phenotype. How are these remodeling events superimposed onto the regulatory state of a differentiated neuron? Previous screens for mutants with an impaired dauer entry phenotype (Daf-d for “dauer defective” or Daf-c for “dauer constitutive”) have identified three hormonal systems involved in this process: an insulin-like signaling pathway, a TGFβ signaling pathway and a nuclear steroid receptor pathway (**Fig.1A**) (Antebi et al., 2000; Fielenbach and Antebi, 2008; Kimura et al., 1997; Lin et al., 1997; Ogg et al., 1997; Ren et al., 1996). While these pathways have been studied to a varying extent in different tissue types, it remains to be better understood where and when these systems act to remodel individual tissue types, specifically in regard to the remodeling of the nervous system.

**Fig.1.**
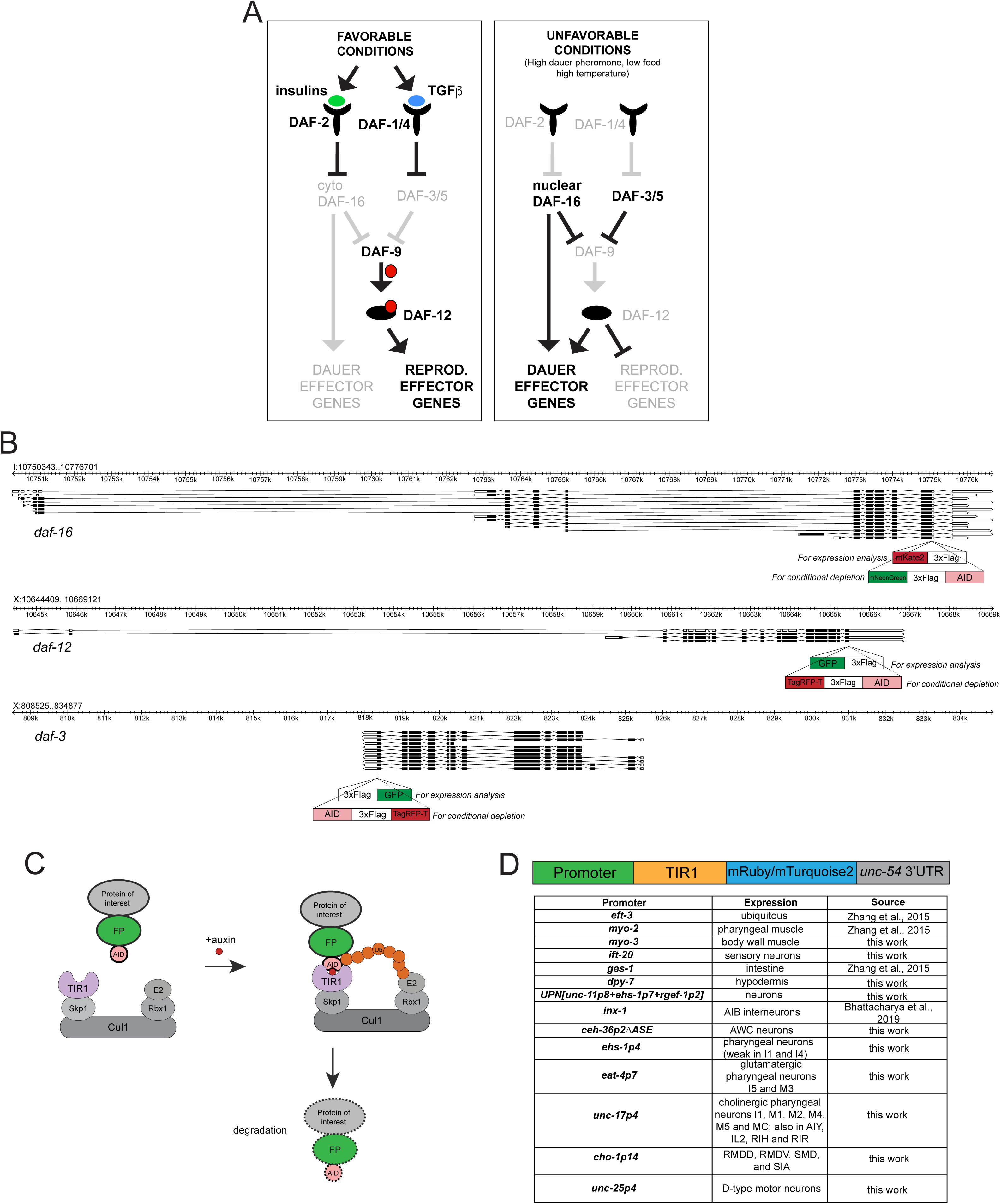
**Reagents generated for this study** (A) Overview of the dauer pathways, with DAF-3/SMAD, DAF-16/FoxO and DAF-12/VDR as transcriptional effectors. (B) Genomic loci of *daf-16, daf-12* and *daf-3*. The insertions sites and schematics of the expression reporters and of the AID tag are shown. (C) Schematic of the AID system. Skp1, Cul1, Rbx1, E2 are phylogenetically conserved components of the E3 ligase complex. TIR1 is a plant-specific substrate recognizing subunit of the E3 ligase complex. Auxin-inducible degron (AID), fused to a protein of interest, is bound by TIR1 in the presence of auxin, which leads to ubiquitination and proteasomal degradation of the protein of interest. (D) Summary of TIR1 transgenes utilized in this study. See Methods for details on TIR constructs.

The focus of action of the TGFβ signaling pathway for dauer formation appears to lie in the nervous system, as assessed by the mosaic analysis of the *daf-4* TGFβ receptor (Inoue and Thomas, 2000) and the rescue of neuronal remodeling events through neuronal expression of *daf-4* (Nolan et al., 2002). The situation is less clear for the insulin-like receptor pathway, whose major effector is the ubiquitously expressed DAF-16/FoxO transcription factor (TF), which translocates into the nucleus upon entry into the dauer stage (Lin et al., 2001). Two studies, using transgenic rescue approaches, have proposed distinct foci of action for *daf-16* in controlling entry into the dauer stage: One study showed that supplying *daf-16* into the nervous system of *daf-16; daf-2* double mutant animals restored their ability to undergo dauer remodeling (Libina et al., 2003), while another study showed that the rescue is only achieved with intestinal expression of *daf-16* in the same genetic background (Hung et al., 2014).

The focus of action of DAF-12 for dauer remodeling events is even less clear (Antebi, 2013; Sims et al., 2016). DAF-12 is a nuclear steroid receptor that responds to *C. elegans* steroids, the dafachronic acids (DA), which represent endogenous signals of the feeding state of the animal (Motola et al., 2006). DAF-12 and its paralogue NHR-8 are similar to the vertebrate xenobiotic nuclear receptors PXR and CAR and to the VDR protein, a nuclear receptor for vitamin D, an endogenously produced neurosteroid (Cui et al., 2017). Intestinally expressed NHR-8, a likely PXR and CAR ortholog (Lindblom et al., 2001; Timsit and Negishi, 2007), responds to xenobiotic substances, while DAF-12, like the vertebrate VDR, responds to endogenously produced steroids. Similar to DAF-12, the vertebrate VDR is broadly expressed in the brain and mediates the many effects that vitamin D has on neuronal development and plasticity (Cui et al., 2017). The focus of action of VDR is not well understood, and the same holds for DAF-12, as far as any tissue-remodeling event during dauer entry is concerned.

In this paper, we address a number of presently unresolved questions about these three signaling systems. Do they operate autonomously within target tissues that undergo cellular remodeling? Or do they serve to relay internal signals to non-autonomously control cellular remodeling? Do they operate together within specific cellular context, or do they work sequentially? Do they only control the initial remodeling events upon dauer entry or are they continuously required to maintain the remodeled state? To address these questions, we generated conditional and fluorescently labeled alleles of the effectors of each of the three hormonal signaling systems, the SMAD TF DAF-3, the FoxO TF DAF-16 and the steroid receptor DAF-12. We show that each of the three TFs displays distinctive dynamics in expression and localization throughout different cell types during continuous development and dauer entry. Temporally controlled removal shows their importance not only initiating but also maintaining the remodeled state. Tissue and cell-type specific removal of each of these TFs reveals complex requirements for these systems and shows that one hormonal axis acts mostly cell-autonomously, while two others act in both target cells (i.e. cell-autonomously), but also act outside the remodeled organ. Such cell non-autonomous activities reveal a number of interorgan signaling axes, from the nervous system to the gut and muscle and also from the gut to the nervous system. We show that these hormonal signaling systems cooperate with hardwired terminal selector type transcription factors to control cellular remodeling. Terminal selectors provide the cellular specificity and potential for remodeling, while these signal-dependent TFs either promote or inhibit the ability of terminal selectors to control remodeling events.

## RESULTS

### Expression pattern of *daf-3/SMAD, daf-12/VDR* and *daf-16/FoxO*

The expression patterns of *daf-3/SMAD*, *daf-16/FoxO* and *daf-12/VDR,* the transcriptional effectors of the three hormonal signaling systems (**Fig.1A**) have previously been examined with multicopy reporter transgenes (Antebi et al., 2000; Lee et al., 2001; Ogg et al., 1997; Patterson et al., 1997). To avoid potential issues with reporter transgenes (such as overexpression or lack of *cis*-regulatory control elements), we tagged the endogenous *daf-3/SMAD*, *daf-16/FoxO* and *daf-12/VDR* loci with different fluorescent tags using CRISPR/Cas9-mediated genome engineering (**Fig.1B**). For all three genes, we chose C-terminal fusions, since they tag all isoforms of the respective loci (**Fig.1B**). The analysis of the three endogenously tagged reporter alleles largely corroborated previously described sites of expression, based on reporter constructs, but also revealed novel spatiotemporal dynamics of gene expression.

The *daf-3* reporter alleles *ot875* and *ot877* generally have very low expression levels (**Fig.2A**). The expression is most prominent in L1 larvae, where it is observed in all tissue types, whereas it is downregulated in subsequent larval stages and in adults, being only detectable in neurons and the intestine. Starvation upregulates *daf-3::GFP* levels not only in L1 larvae, but also in later larval stages and adults (**Fig.2A; Suppl.Fig.S2**). In dauers, expression levels of *daf-3::GFP* are lowest, with only nuclei of some head neurons visible.

**Fig.2.**
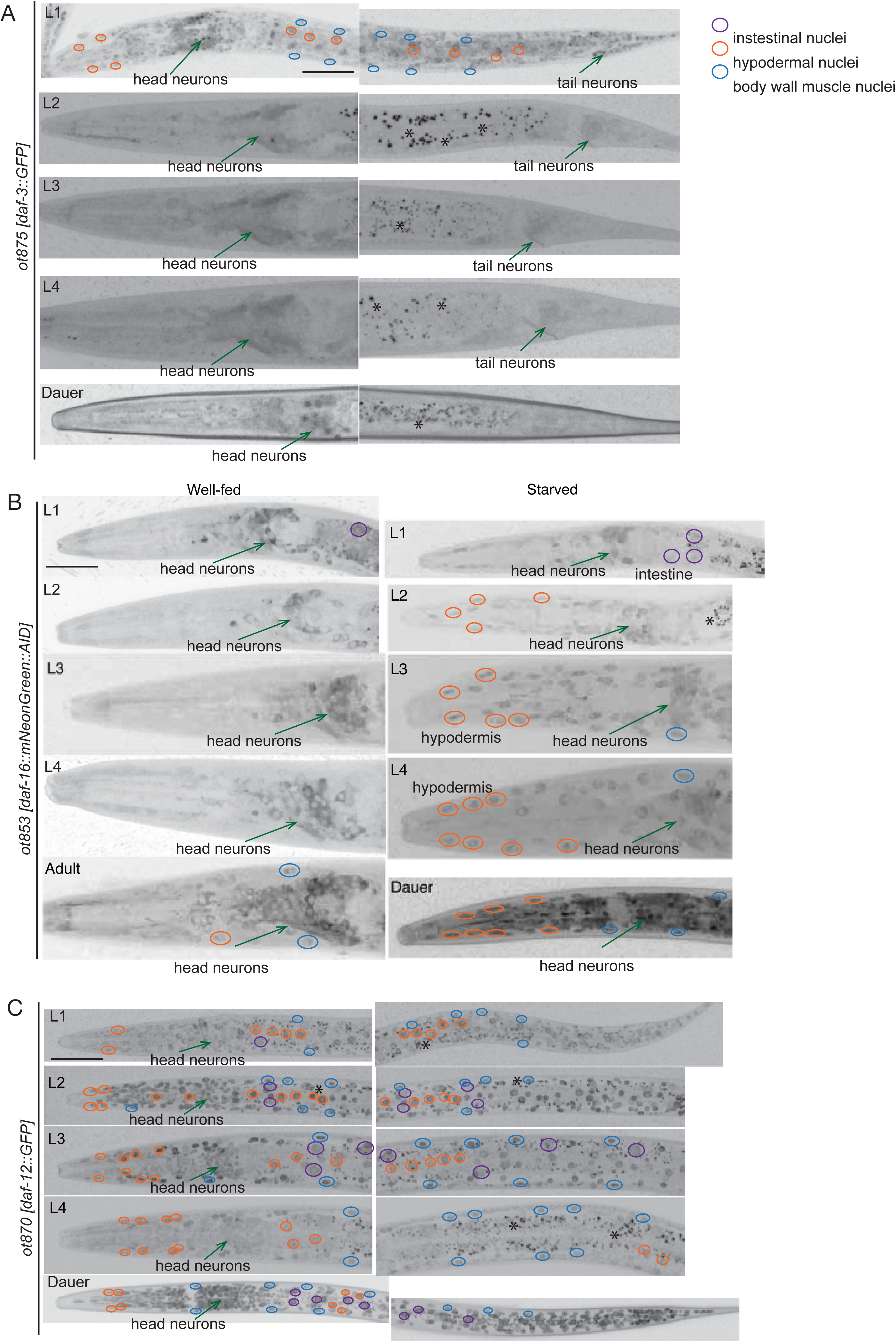
**Expression pattern of *daf-16/FoxO*, *daf-12/VDR* and *daf-3/SMAD*** (A) Expression of the *daf-3::GFP* CRISPR allele at different stages in development (in well-fed conditions). Anterior is to the left on all images. Scale bar, 20 µm (same for all images). (B) Expression of the *daf-16::mKate2* CRISPR allele at different stages in development and nutritional states. Different tissue types are indicated with color-coded circles (or an arrow, in the case of neurons). (C) Expression of the *daf-12::GFP* CRISPR allele at different stages in development (in well-fed conditions).

In well-fed larvae and adults, the *daf-16* CRISPR reporter alleles *ot821* and *ot853* are expressed in the cytoplasm of neurons, most prominently, in the sensory neurons of the head (**Fig.2B**). Non-neuronal tissues show only weak expression. After food deprivation, the tagged DAF-16 protein is upregulated in non-neuronal tissues, where it is mostly nuclear (**Fig.2B**). However, in neurons of starved larvae and adults, DAF-16 remains mostly cytoplasmic. In striking contrast, in dauers, the tagged DAF-16 protein displays nuclear localization throughout all tissues, including neurons (**Fig.2B**).

The *daf-12* reporter alleles *ot870* and *ot874* displayed ubiquitous expression, with highest expression levels during reproductive development observed at L2 (**Fig.2C**), largely correlating with the timing of *daf-12’s* heterochronic gene activity (Antebi et al., 1998). While starvation at different stages does not appear to impinge on the expression level of *daf-12, daf-12* expression is significantly stronger throughout all tissues in the dauer stage (**Fig.2C**) (contrasting earlier reports using a multicopy, cosmid-based reporter array of *daf-12*; (Antebi et al., 2000)).

### DAF-3/SMAD, DAF-12/VDR and DAF-16/FoxO are continuously required to maintain the dauer stage

To deplete gene function in a spatially and temporally controlled manner, we generated conditional alleles by inserting an auxin-inducible degron (AID) into each of the loci (Zhang et al., 2015)(**Fig.1B,C**). This provided a genetic strategy that is orthogonal to the cell-specific rescue approaches previously used to assess the focus of *daf-16/FoxO* gene function (Hung et al., 2014; Libina et al., 2003; Wolkow et al., 2000). For *daf-3/SMAD* and *daf-12/VDR*, no studies on the cellular focus of action for the dauer decision have been conducted so far.

We found that tagging each of the TF loci had no detectable effect on gene function in the context of dauer formation, as assessed by crossing the tagged loci into distinct Daf-c mutant backgrounds, whose Daf-c phenotype is suppressed by reduction of *daf-3/SMAD*, *daf-12/VDR* or *daf-16/FoxO* gene function (described further below). No suppression effects were observed with the tagged loci for any of the three TFs.

To achieve spatio-temporally controlled gene depletion, we generated transgenic lines that express the substrate-recognizing subunit of the plant ubiquitin ligase, TIR1, an essential component of the AID system (Zhang et al., 2015), in a number of distinct tissue types. In addition to previously generated TIR1 transgenes (ubiquitously expressed *ieSi57*, pharyngeal muscle-expressed *ieSi60* and intestine-expressed *ieSi61*) (Zhang et al., 2015), we generated new transgenes that express TIR1 in tissue- or cell-specific manners (described in Methods and summarized in **Fig.1D**). In the presence of auxin, these TIR1 transgenes effectively remove DAF-3/SMAD::TagRFP-T::AID, DAF-12::TagRFP-T::AID and DAF-16::mNeonGreen::AID protein from individual cell types, as assessed by the loss of fluorescent signals (shown in **Suppl. Fig.S1** and other main Figures discussed later). To corroborate efficient gene depletion at a functional level, we tested whether AID-mediated protein removal, using ubiquitously expressed TIR1, can recapitulate the *daf-3/SMAD*, *daf-12/VDR* and *daf-16/FoxO* null mutant phenotypes. To this end, we crossed the tagged loci into the *daf-7(e1372)* or *daf-2(e1370)* Daf-c mutant backgrounds that were previously shown to be suppressed by *daf-3/SMAD*, *daf-12/VDR* or *daf-16/FoxO* null mutants, respectively (Gottlieb and Ruvkun, 1994; Kenyon et al., 1993; Thomas et al., 1993). Eggs of these strains were plated on control or auxin-treated plates and allowed to grow for three days at 25°C (**Fig.3A**). Without ubiquitous TIR1 expression and/or without auxin addition, neither of the tagged loci suppressed the respective Daf-c mutant background, demonstrating that the reporter tagging does not affect gene function (**Fig.3B, Fig.4A-C**). However, the combination of ubiquitous TIR1 expression and auxin addition resulted in a full suppression of the respective Daf-c mutant phenotypes, demonstrating the effectiveness of the AID system (**Fig.3B**).

**Fig.3.**
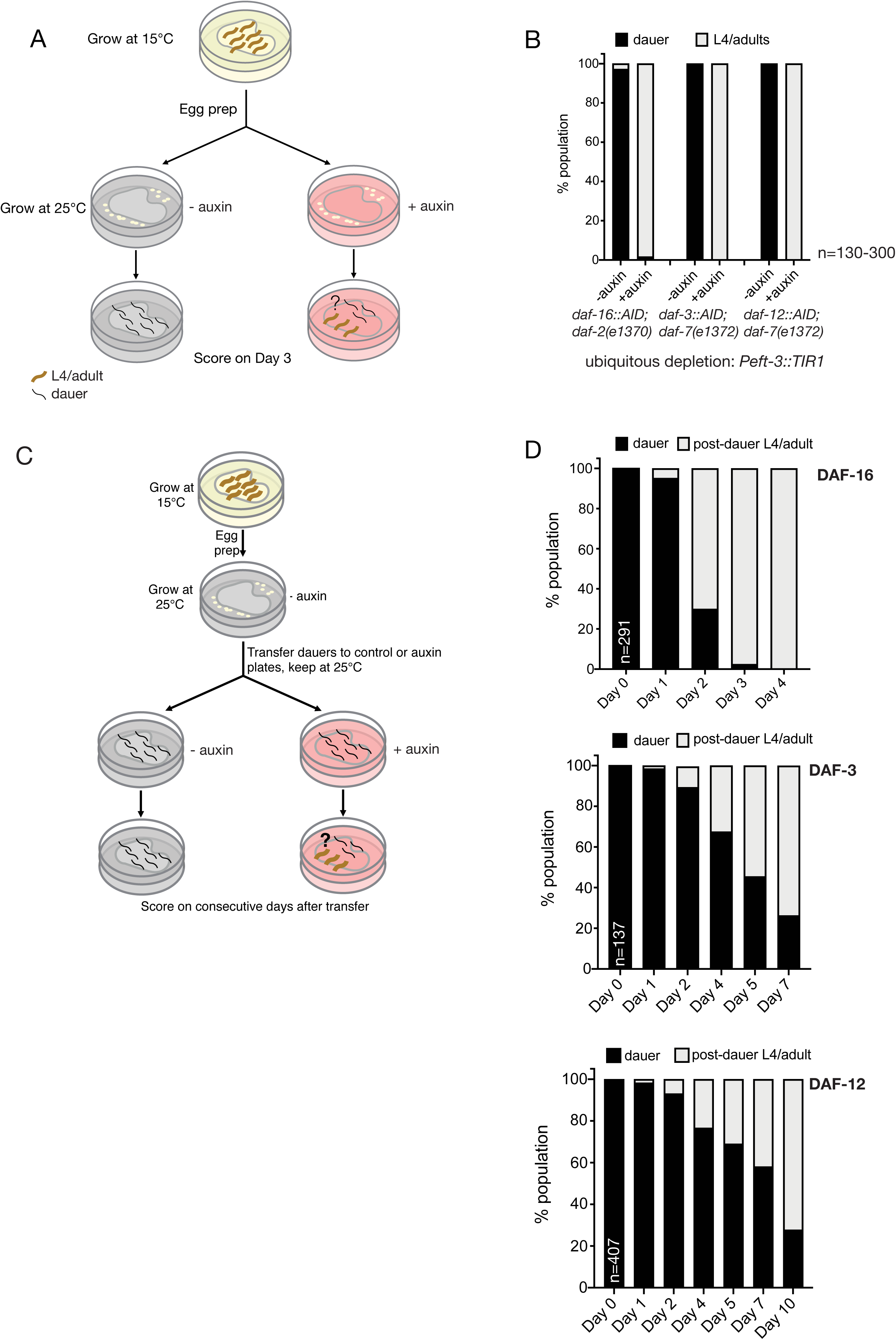
**Requirement of DAF-16/FoxO, DAF-12/VDR and DAF-3/SMAD for initiation and maintenance of the dauer state** (A) Schematic of the experimental design for testing conditional alleles of *daf-16*, *daf-12* and *daf-3* for the requirement of the respective proteins in initiation of dauer formation (also applies to experiments described in later figures). (B) Dauer formation is suppressed upon ubiquitous depletion of DAF-16/FoxO, DAF-12/VDR and DAF-3/SMAD, which serves as a positive control for subsequent experiments with tissue-specific protein depletion. (C) Schematic of the experimental design for testing conditional alleles of *daf-16*, *daf-12* and *daf-3* for the requirement of the respective proteins in maintenance of the dauer state. (D) Upon ubiquitous depletion of DAF-16/FoxO, DAF-12/VDR and DAF-3/SMAD after dauer formation, worms exit the dauer state and initiate post-dauer development, indicating the requirement of the three proteins in the active maintenance of the dauer state.

**Fig.4.**
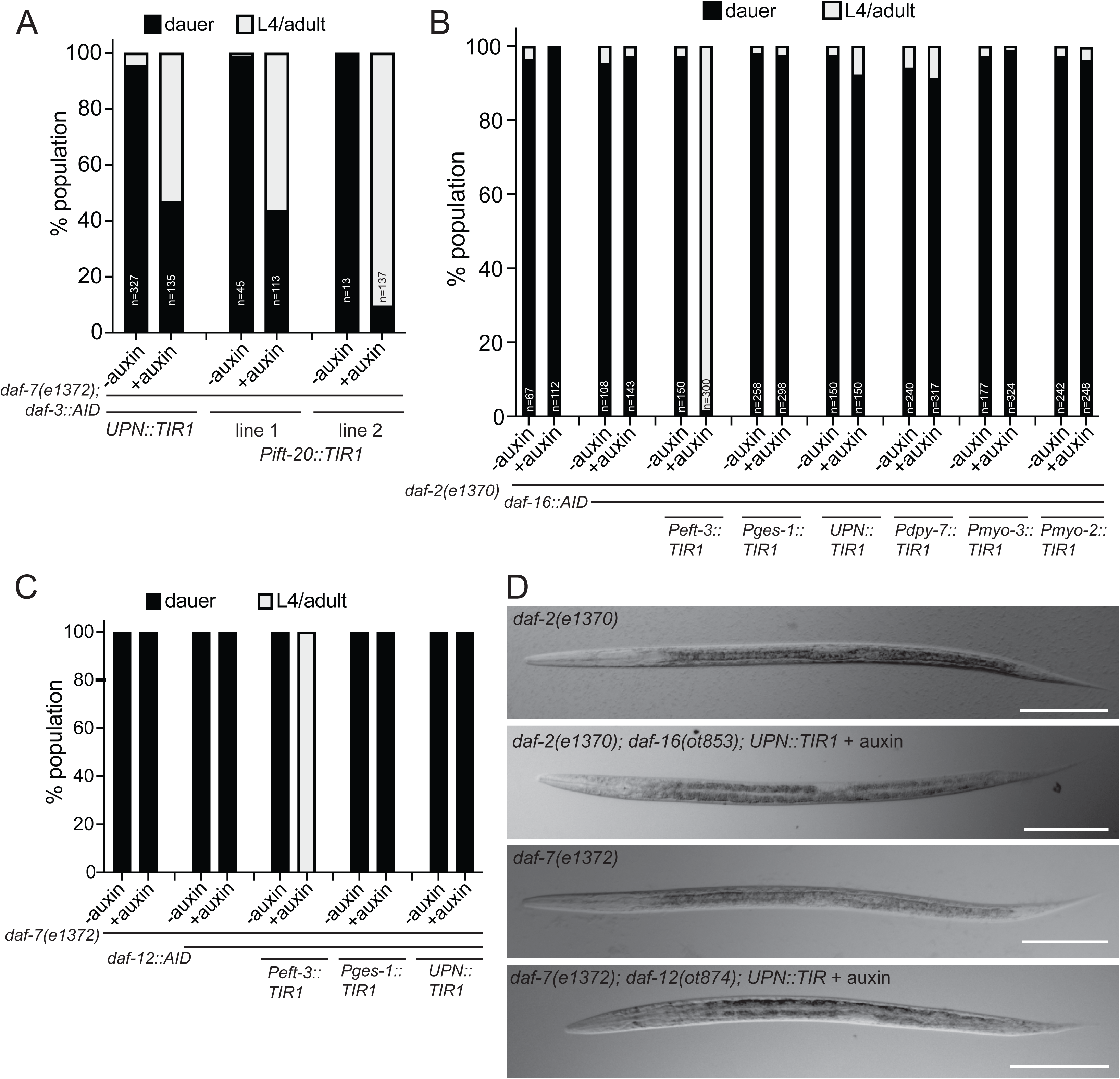
**Neuronal depletion of DAF-3/SMAD, but not of DAF-16/FoxO or DAF-12/VDR, suppresses dauer formation** (A) Quantification of dauer formation upon depletion of DAF-3/SMAD with a panneuronal (*otIs730[UPN::TIR1::mTurquoise2]*) and a pansensory TIR1 (*otEx[Pift-20::TIR1::mRuby]*) drivers. (B) Quantification of dauer formation upon tissue-specific depletion of DAF-16/FoxO. (C) Quantification of dauer formation upon tissue-specific depletion of DAF-12/VDR. (D) Overall appearance of dauers with panneuronal depletion of DAF-16/FoxO and DAF-12/VDR is superficially similar to that of the background Daf-c strains, *daf-2(e1370)* and *daf-7(e1372)*, respectively. Scale bars: 60 μm.

Next, we asked whether these three TFs are only required for the commitment to the dauer remodeling program or whether they are continuously required to maintain the remodeled state. To address this question, we performed auxin shift experiments, in which eggs are initially plated on control plates and allowed to develop into dauers at 25°C, which are then transferred to auxin plates on day 3 (or to auxin(-) plates for the control animals) (**Fig.3C**). Depletion of DAF-16/FoxO, DAF-3/SMAD or DAF-12/VDR after dauer formation results in dauer exit, indicating that all three proteins are required for the maintenance of the dauer state (**Fig.3D**).

### Neuronal removal of DAF-3/SMAD, but not of DAF-12/VDR or DAF-16/FoxO, suppresses dauer formation

After assessing temporal requirements, we sought to define the focus of action of the three hormonal systems, through cell-type specific gene depletion. Previous mosaic analysis of the TGFβ receptor *daf-4* indicated that TGFβ signaling is required in the nervous system to control dauer formation (Inoue and Thomas, 2000). In accordance with these findings, we find that panneuronal depletion of DAF-3/SMAD, using a panneuronal (UPN) TIR1 driver line, is indeed able to significantly suppress the DAF-3/SMAD-dependent Daf-c phenotype of *daf-7(e1372)* mutants. To further delineate the focus of DAF-3/SMAD action, we removed DAF-3/SMAD exclusively from ciliated sensory neurons, using an *ift-20::TIR1* transgenic line and found that this also significantly suppresses the Daf-c phenotype of *daf-7(e1372)* animals (**Fig.4A**). We conclude that DAF-3/SMAD acts in sensory neurons to non-autonomously control remodeling of multiple tissue types during dauer remodeling. This is consistent with previous transgenic rescue assays of mutant animals in which the TGFβ receptor DAF-4 was mutated, which suggested a possible function in sensory neurons in the context of controlling the expression of sensory receptor proteins (Nolan et al., 2002).

Previously published transgenic overexpression approaches indicated that *daf-16/FoxO* function in either the nervous system or the gut is sufficient to rescue *daf-16/FoxO* defects (Hung et al., 2014; Libina et al., 2003), but these studies did not address in which tissue type *daf-16/FoxO* is normally required to promote dauer remodeling. Using the same panneuronal TIR1 driver line that functionally removed DAF-3/SMAD from the nervous system to suppress dauer formation, we found that panneuronal removal of DAF-16/FoxO was not able to suppress the Daf-c phenotype of *daf-2(e1370)* mutants (**Fig.4B**). Similarly, panneuronal removal of DAF-12/VDR did not suppress the Daf-c phenotype of *daf-7(e1372)* mutants (**Fig.4C**). In both cases, dauer appear morphologically like normal dauers and they also retain their chemical resistance to SDS, a key feature of dauers (**Suppl.Fig.S3**)(Cassada and Russell, 1975). Also, in both cases, fluorescence microscopy shows that the panneuronal TIR1 driver effectively removed DAF-16::mNG::AID and DAF-12::TagRFP-T::AID protein from the nervous system (**Suppl. Fig.S1**).

An alternative focus of action for *daf-16/FoxO* has been proposed through transgenic rescue experiments, which showed that re-introduction of *daf-16/FoxO* exclusively in the intestine was able to revert the dauer suppression phenotype of *daf-2(e1370); daf-16(Df50)* double mutants (Hung et al., 2014). However, our own examination of this previously published strain revealed that the suppression was not complete, namely that the transgenic dauers fail to remodel their pharynx and continue to pump, unlike *daf-2(e1370)* dauers, although they retain SDS-resistance, a distinctive characteristic of dauers (**Suppl. Fig.S4**). To address this issue from an orthogonal angle, we examined whether intestinal depletion of DAF-16/FoxO is able to suppress the Daf-c phenotype of *daf-2(e1370)* mutants. We observed no suppression (**Fig.4B**), arguing that DAF-16/FoxO is at least not solely required in the intestine to promote entry into the dauer stage. We also detected no effect of DAF-12/VDR removal from the intestine on the Daf-c phenotype of *daf-7(e1372)* mutants (**Fig.4C**).

### Neuronal removal of DAF-16/FoxO affects dauer-specific sensory receptor expression changes

Our observation that animals that lack DAF-16/FoxO or DAF-12/VDR activity in the nervous system can still enter the dauer stage provided us with the opportunity to ask what functions DAF-16/FoxO and DAF-12/VDR play within the nervous system of dauer animals for neuronal remodeling. In the next few sections, we will first describe our analysis of DAF-16/FoxO and at the end of this paper we will describe our results with DAF-12/VDR.

One prominent neuronal remodeling event that relates to altered chemosensory behavior of dauers is evidenced by the many expression changes of olfactory-type G-protein coupled receptors upon entry into the dauer stage (Nolan et al., 2002; Peckol et al., 2001; Vidal et al., 2018). For example, we previously reported that the *sri-9* GPCR gene, which is normally expressed only in the ADL neuron class in the head of the worm, becomes induced in 8 additional neuron classes upon entry into the dauer stage (Vidal et al., 2018). We find that panneuronal removal of DAF-16/FoxO largely suppresses the induction of *sri-9* reporter gene expression in these additional neuron classes (**Fig.5A**), suggesting that DAF-16/FoxO functions within the nervous system to control changes in GPCR expression. However, factors other than DAF-16/FoxO (or other than its function in neurons) must also play a role, since we find that the dauer-specific induction of *sra-25* in the ADL neurons (Vidal et al., 2018) still occurs after panneuronal DAF-16/FoxO depletion (**Fig.5B**). The *daf-16*-independence of *sra-25* induction in ADL of dauers may relate to the fact that *sra-25* expression persists even after exit from the dauer stage (Vidal et al., 2018), i.e., after inactivation of DAF-16 in post-dauer animals.

**Fig.5.**
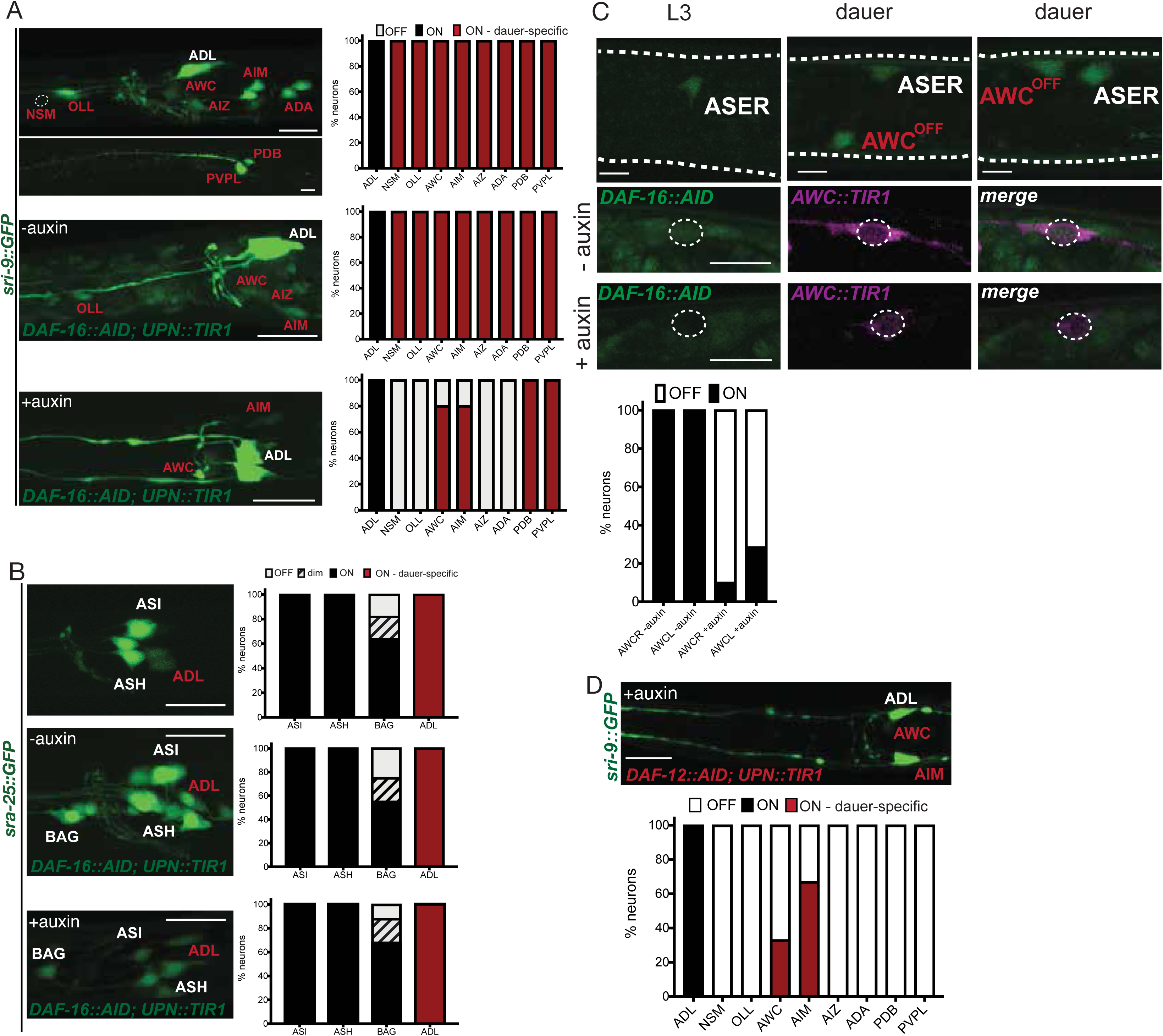
**Neuronal removal of DAF-16/FoxO and DAF-12/VDR affects dauer-specific sensory receptor expression changes.** (A) *sri-9::GFP* (*otIs732*) gains expression in multiple neurons in dauer (red). The dauer-specific expression is largely dependent on the neuronal function of DAF-16/FoxO. Left, head images of control and auxin-treated dauers with panneuronal depletion of DAF-16/FoxO. Right, quantification of *sri-9::GFP* expression in each neuronal type. n=22 (control) and n=10 (auxin-treated). Scale bars, 10 µm. (B) *sra-25::GFP* (*sIs12199*) gains expression in ADL neurons in dauer (red). The dauer-specific expression is not dependent on neuronal function of DAF-16/FoxO. Left, head images of control and auxin-treated dauers with panneuronal depletion of DAF-16/FoxO. Right, quantification of *sra-25::GFP* expression. n=22 (control) and n=10 (auxin-treated). Scale bars, 10 µm. (C) *gcy-5::NLS::GFP* (*otIs586*) expression is gained in AWC^OFF^ neuron in dauer. Above, images of AWC in control and auxin-treated dauers with panneuronal depletion of DAF-16/FoxO. Below, quantification of *gcy-5::NLS::GFP* expression. n=12. Scale bars, 5 µm. (D) The dauer-specific expression of *sri-9::GFP* (*otIs732*) also depends on the neuronal function of DAF-12/VDR. Above, image of an auxin-treated dauer with panneuronal depletion of DAF-12/VDR. Below, quantification of *sri-9::GFP* expression in each neuronal type. Scale bar, 10 µm.

To extend our probing of the impact of DAF-16/FoxO on sensory receptor expression and to address the question of cell autonomy with more precise cellular resolution, we turned to another sensory receptor system, the receptor type guanylyl cyclases (rGCs). We have previously studied this family of genes in the context of taste perception in non-dauer stage animals (Ortiz et al., 2009). We noted that expression of one rGC, *gcy-5*, which is normally exclusively expressed in the ASEL salt receptor neuron (Yu et al., 1997), becomes activated in the AWC^OFF^ olfactory neuron during entry into the dauer stage (**Fig.5C**). We established an AWC-specific TIR1 driver line to ask whether DAF-16/FoxO acts specifically in AWC neurons to promote *gcy-5* expression during dauer remodeling, and indeed found this to be the case (**Fig.5C**).

### Cell autonomous control of electrical synapse remodeling by DAF-16/FoxO

Another prominent set of remodeling events that we recently described to occur upon dauer entry are changes in expression of electrical synapse proteins, the innexins (Bhattacharya et al., 2019). For example, expression of the *inx-6* innexin becomes exclusively induced in the AIB interneurons of dauers. We had previously shown that this induction requires DAF-16/FoxO activity in the AIB interneurons (Bhattacharya et al., 2019). We extended these previous findings by using the example of AIB neurons as a test case to ask whether this cell-autonomous process requires continuous DAF-16/FoxO expression. Alternatively, DAF-16/FoxO may only be transiently required, for example, as a pioneer factor to permit the induction of *inx-6* expression. By auxin-shifting the dauers initially grown in control conditions, we found that DAF-16/FoxO is continuously required to maintain *inx-6* during the dauer stage (**Fig.6A**), which is consistent with the continuous requirement for DAF-16/FoxO to retain animals in the dauer stage, as described above (**Fig.3C, D**).

**Fig.6.**
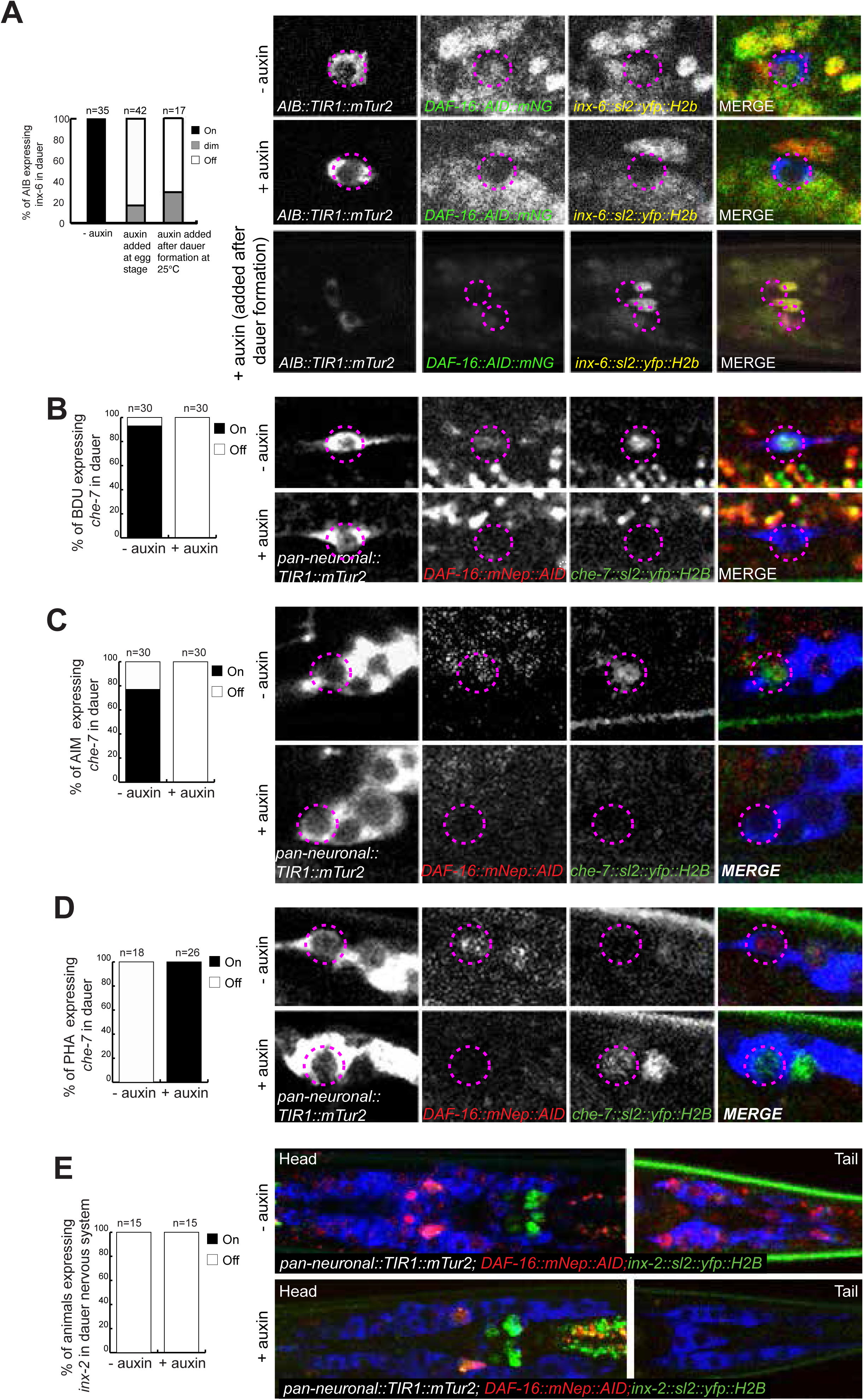
**Neuronal removal of DAF-16/FoxO largely controls dauer-specific expression changes of electrical synapse components.** (A) The dauer-specific expression gain of an *inx-6* reporter allele *(ot804)* in AIB is affected upon continuous AIB-specific depletion of DAF-16/FoxO, as well as in animals where DAF-16/FoxO is depleted in AIB after dauer remodeling. (B) A *che-7* reporter (*otEx7112)* expression is gained in BDU neurons in dauer. This dauer-specific *che-7* expression is lost upon panneuronal depletion of DAF-16/FoxO in auxin-treated dauers. (C) A *che-7* reporter (*otEx7112)* expression is gained in AIM neurons in dauer. This dauer-specific *che-7* expression is lost upon panneuronal depletion of DAF-16/FoxO in auxin-treated dauers. (D) A *che-7* reporter (*otEx7112)* expression is downregulated in PHA neurons in dauer. This dauer-specific downregulation of *che-7* expression is affected upon panneuronal depletion of DAF-16/FoxO in auxin-treated dauers. (E) Expression of an *inx-2* reporter allele (*ot906)* is downregulated in multiple neurons in dauer. This dauer-specific downregulation of *inx-6* expression is unaffected upon panneuronal depletion of DAF-16/FoxO in auxin-treated dauers. In all the animals scored, AID-tagged DAF-16/FoxO was also completely depleted in the presence of auxin.

We extended our analysis to a number of additional innexin genes, particularly also considering innexin genes whose expression does not become upregulated, but rather becomes downregulated upon entry into the dauer stage. One such example is *che-7*, an innexin that is upregulated in some neurons, but is downregulated in other neuron types upon dauer entry (Bhattacharya et al., 2019). We find that the upregulation of the *che-7* innexin in a number of neuron types (including BDU, NSM, and AIM) fails to occur after panneuronal depletion of DAF-16, consistent with its cell-autonomous activity (**Fig.6B,C and Suppl. Fig.S5**). Intriguingly, the dauer-specific downregulation of *che-7* in the PHA and OLL neurons is also abolished upon panneuronal DAF-16 depletion (**Fig.6D**), indicating that DAF-16, either directly or indirectly, activates or represses genes depending on cellular context.

However, DAF-16 is not required for the regulatory plasticity of every single innexin gene. We find that panneuronal DAF-16 is not required for the downregulation of *inx-2*, which occurs in multiple neurons types upon dauer entry (**Fig.6E**).

### Remodeling of locomotory behavior requires neuronal DAF-16/FoxO

Entry into the dauer stage is accompanied by remarkable changes in locomotory behavior (Bhattacharya et al., 2019; Cassada and Russell, 1975; Gaglia and Kenyon, 2009). Among such changes are a decrease in the wave amplitude of the sinusoidal movement patterns of the animals and a decrease in various aspects of reversal behavior, including omega turn behavior, as quantified using a semi-automated WormTracker system (Bhattacharya et al., 2019). We used the same system to quantify locomotor aspects of dauer-stage animals in which DAF-16/FoxO was panneuronally depleted. Strikingly, these animals retain many patterns of non-dauer locomotion, i.e., they fail to undergo the alterations normally associated with dauer behavior. For example, among other locomotory features, panneuronally DAF-16/FoxO-depleted dauer animals display an increased amplitude of sinusoidal movement and an increase in omega turn frequency (**Fig.7A, B**). These animals move much faster than vehicle controls as well as *daf--2(e1370)* dauers, and pause less during locomotion. Comparison with fed L3 larvae of the *daf-2(e1370)* background strain show that panneuronally DAF-16-depleted dauers behave more L3-like than bona fide dauers (**Fig.7A, B**). Depleting DAF-16 from body wall muscle did not result in any significant changes in locomotory behavior (**Suppl. Fig.S6**), but, as we will describe further below, it did have other effects on the animal.

**Fig.7.**
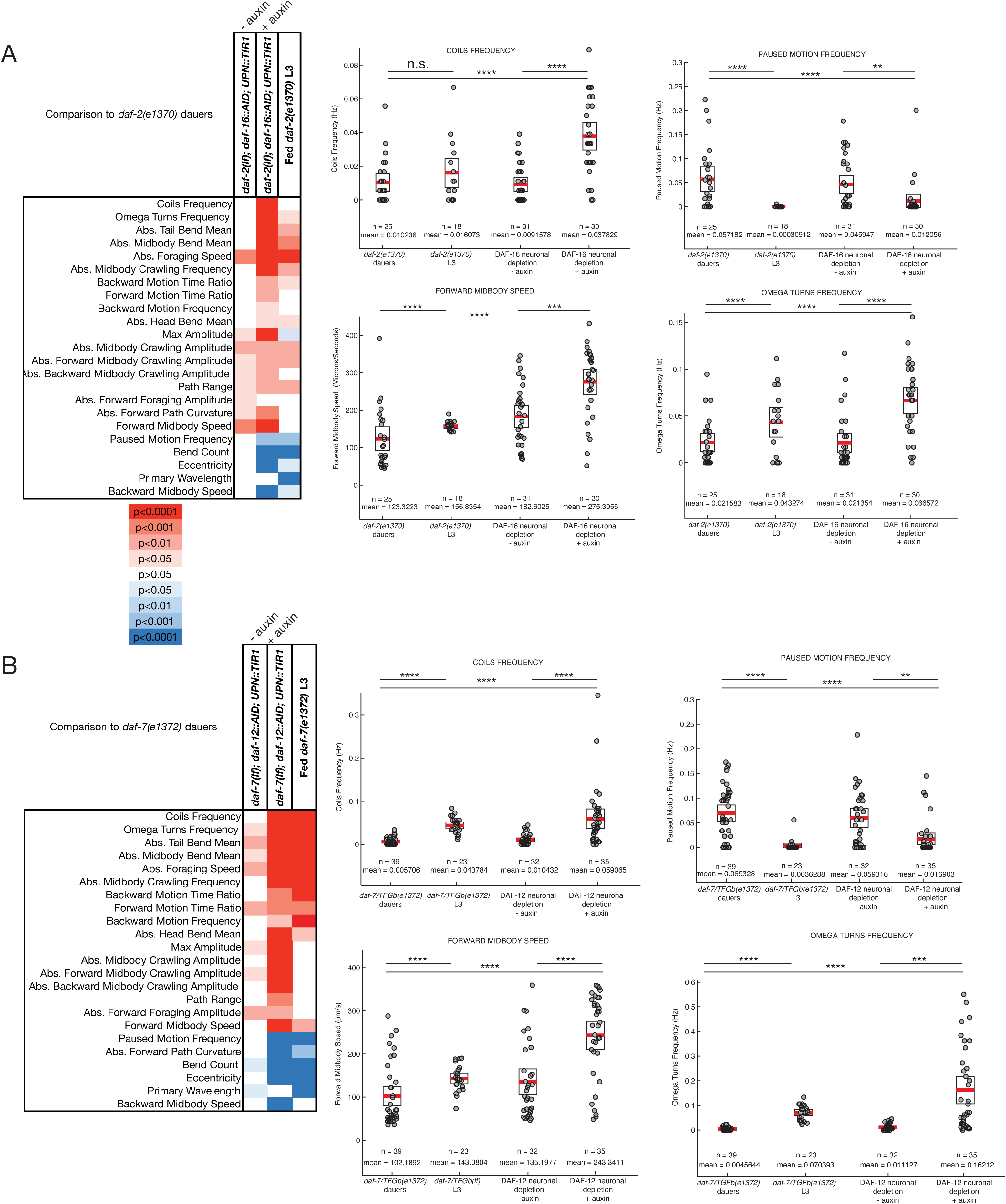
**DAF-16/FoxO and DAF-12/VDR act in the nervous system to affect dauer-specific locomotory behavior** (A) A heat map of *p*-values associated with locomotory features of panneuronally DAF-16/FoxO-depleted dauers. (B) Representative locomotory features, which distinguish dauers with panneuronal depletion of DAF-16/FoxO from controls. (C) A heat map of *p*-values associated with locomotory features of panneuronally DAF-12/VDR-depleted dauers. (D) Representative locomotory features which distinguish dauers with panneuronal depletion of DAF-12/VDR from controls. Each circle represents the experimental mean of a single animal. Red lines indicate the mean of means and rectangles indicate SEM. Wilcoxon rank-sum tests and false-discovery rate q values for each comparison: n.s., nonsignificant, \**q* < 0.05, \*\**q* < 0.01, \*\*\**q* < 0.001, \*\*\*\**q* < 0.0001.

### Remodeling of the pharynx reveals cell autonomous and non-autonomous function of DAF-16/FoxO

Another conspicuous behavioral change observed upon entry into the dauer stage is the cessation of pharyngeal pumping behavior (Cassada and Russell, 1975). Pharyngeal pumping is controlled by the pharyngeal nervous system, composed of 14 neuron classes that are heavily interconnected and that innervate pharyngeal musculature (Albertson and Thomson, 1976; Avery and Thomas, 1997; Cook et al., 2020). The cessation of pharyngeal pumping during dauer remodeling is also accompanied by an autophagy-dependent shrinkage of pharyngeal muscle tissue (Melendez et al., 2003), thereby resulting in a constricted appearance of the dauer pharynx. We find that panneuronal depletion of DAF-16/FoxO results in a derepression of pharyngeal pumping in dauer animals (**Fig.8A**), indicating that DAF-16/FoxO is normally required in neurons to suppress pumping. Unexpectedly, neuronal depletion of DAF-16/FoxO also results in a failure to constrict pharyngeal musculature (**Fig.8B, C**). To further explore this observation, we also generated a pharyngeal muscle-specific TIR1 driver line and found that pharyngeal muscle depletion of DAF-16/FoxO likewise resulted in a loss of pharyngeal muscle constriction as well as derepression of pharyngeal pumping during the dauer remodeling. Muscle constriction and pharyngeal pumping are not obligatorily linked because in hypomorphic *daf-16(m26)* mutants, pharyngeal muscle tissue of dauer animals fails to constrict (Vowels and Thomas, 1992), but pharyngeal pumping remains suppressed (**Supp. Fig.7A, B**). Vice versa, wild type dauers recovering on food show pharyngeal pumping within 2-3 hours after food introduction (**Suppl. Fig.7C**), even though their pharynx is still constricted.

**Fig.8.**
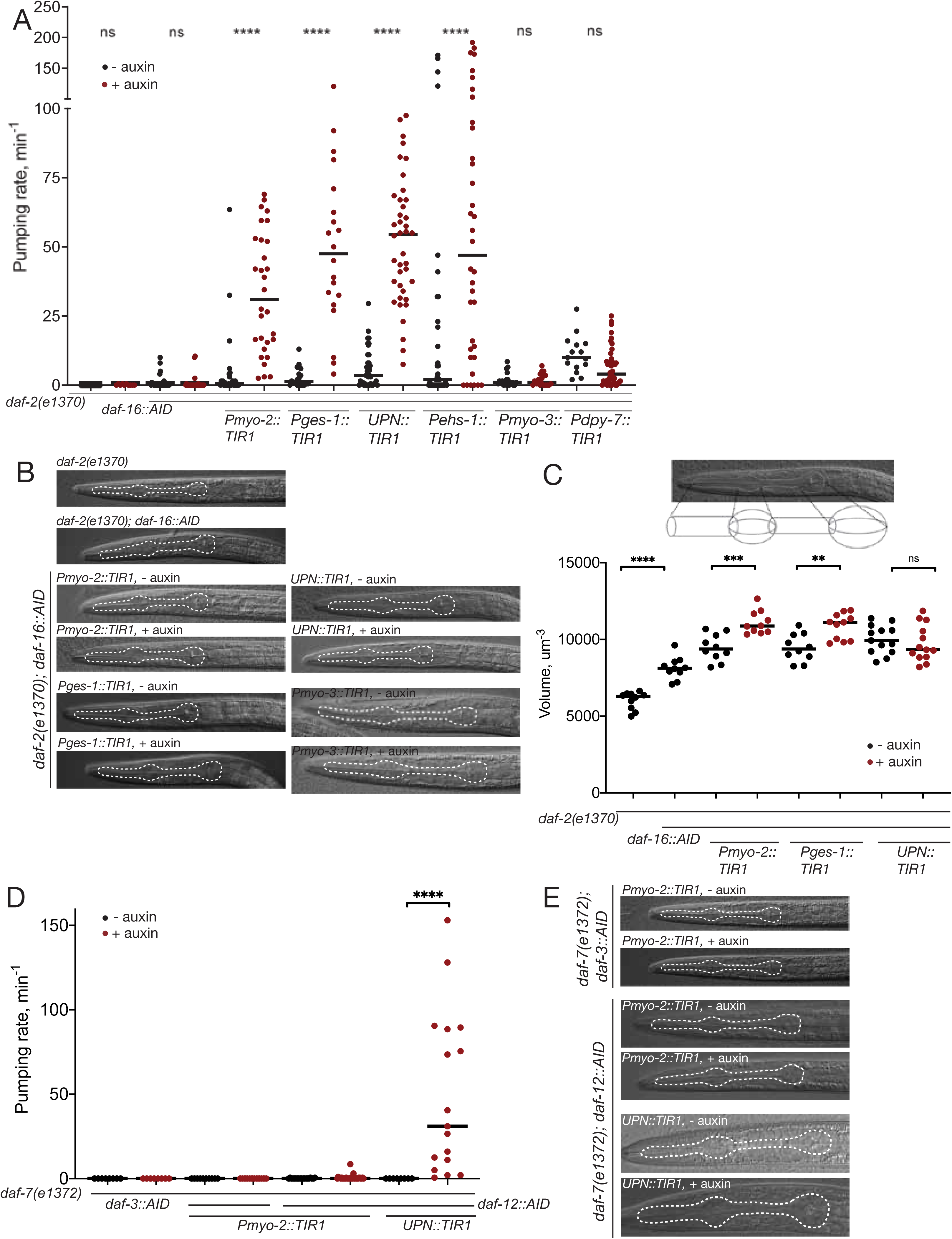
**Control of pharyngeal pumping by DAF-16/FoxO and DAF-12/VDR** (A) Pharyngeal silencing in dauer depends on cell-autonomous and non-autonomous DAF-16 function.Quantification of pharyngeal pumping upon DAF-16/FoxO depletion in pharyngeal muscle (*myo-2*), intestine (*ges-1*), all neurons (*UPN*), pharyngeal neurons (*ehs-1*), body wall muscle (*myo-3*), and hypodermis (*dpy-7*). (B) Pharyngeal constriction depends on cell-autonomous and non-autonomous DAF-16/FoxO function. DIC images of control dauers and dauers with DAF-16/FoxO depleted from pharyngeal muscle, intestine, neurons and body wall muscle. (C) Quantification of changes in pharynx volume upon depletion of DAF-16 from pharyngeal muscle, intestine and neurons. (D) Pharyngeal silencing in dauer depends on the neuronal function of DAF-12. Quantification of pharyngeal pumping upon depletion of DAF-12/VDR in pharyngeal muscle and neurons, as well as upon depletion of DAF-3/Smad in pharyngeal muscle. (E) Pharyngeal constriction in dauer depends on neuronal function of DAF-12/VDR. DIC images of control dauers and dauers with DAF-12/VDR depleted from pharyngeal muscle and neurons as well as with DAF-3/Smad depleted from pharyngeal muscle.

To further dissect the focus of neuronal action of *daf-16,* we depleted DAF-16 exclusively from pharyngeal neurons, using a *cis*-regulatory element from the *ehs-1* gene (Stefanakis et al., 2015) to drive TIR1 expression in pharyngeal, but no other neurons. We found that TIR1 expression driven by this *ehs-1* promoter fragment was strong enough in all the pharyngeal neurons, except I1 and I4, to deplete DAF-16 in the presence of auxin (**Fig.9A, G**). This also resulted in derepression of pharyngeal pumping in dauers (**Fig.9B**). The *ehs-1* promoter fragment used here is also expressed in the head muscle of L1 larvae, but we find that depletion of DAF-16 from all body wall muscles, using the *myo-3* promoter, does not affect pharyngeal activity of dauers (**Fig.8A**). Hence, the effect on pharyngeal pumping seen with the *ehs-1* promoter fragment can be fully attributed to pharyngeal neurons themselves. To further pinpoint the focus of action of DAF-16 in the pharyngeal nervous system, we removed DAF-16 from specific subpopulations of pharyngeal neurons, using driver lines that express TIR1 in either the I1, M1, M2, M4, M5 and MC cholinergic neurons of the pharynx (*unc-17prom4* driver) or in I5 and M3 glutamatergic neurons of the pharynx (*eat-4prom7* driver) (**Fig.9G**)(Serrano-Saiz et al., 2020). We confirmed DAF-16 removal from these pharyngeal neurons by imaging the loss of the fluorescent protein signal of DAF-16::mNG::AID (**Fig.9C, E**). We find that removal of DAF-16 from either neuronal population partially derepressed pumping (**Fig.9D, F**). Since DAF-16 was depleted from non-overlapping neurons in these two TIR1 driver lines (**Fig.9G**), this suggests that DAF-16 function is distributed over different pharyngeal neuron types to silence pumping of the pharynx in the dauer stage.

**Fig.9.**
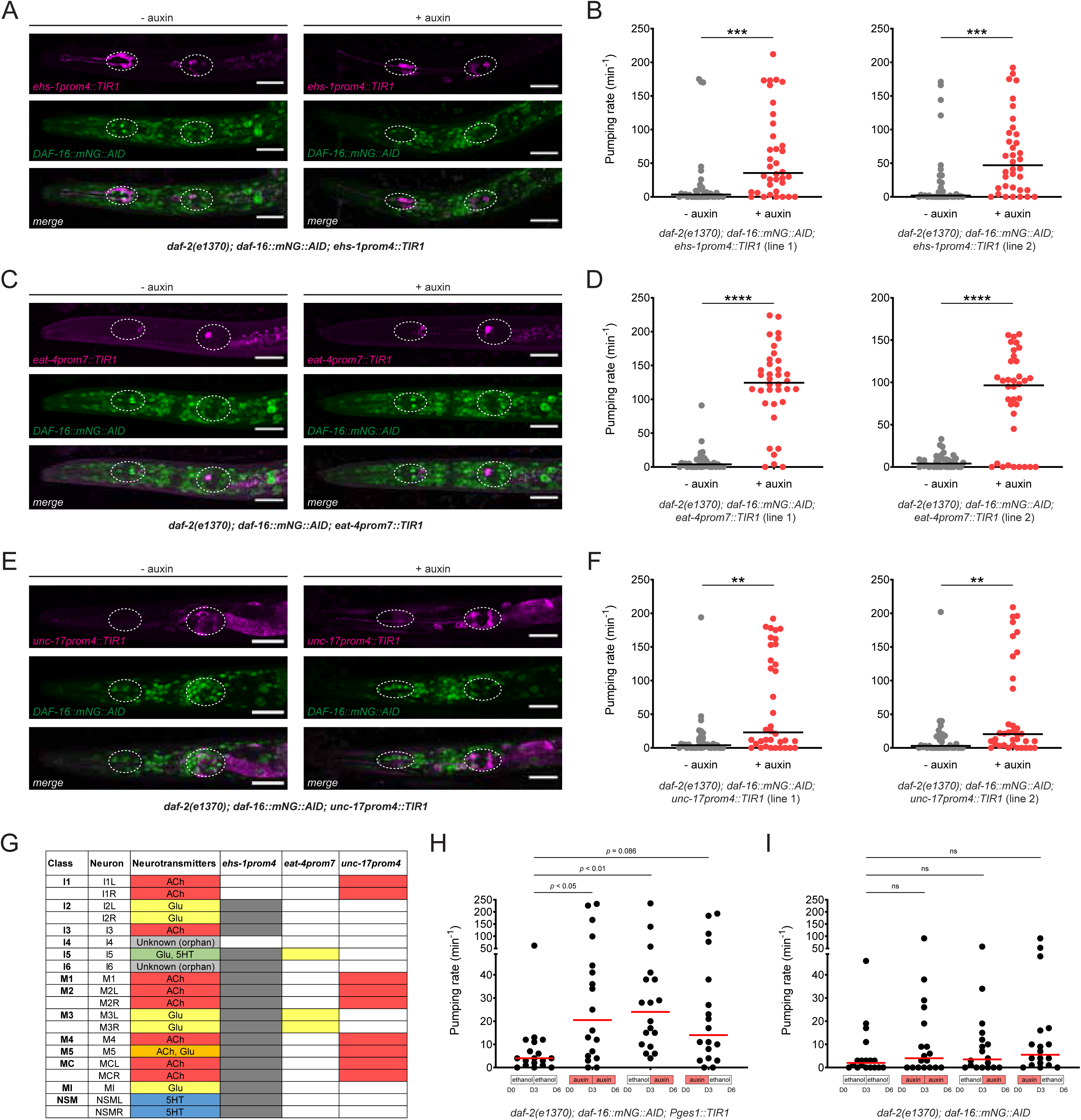
**DAF-16/FoxO activity is distributed across multiple pharyngeal neurons to silence pharyngeal pumping in the dauer stage** (A) DAF-16/FoxO depletion in pharyngeal neurons of *daf-2(e1370)* dauers using *ehs-1prom4*-driven TIR1. In 20/20 animals observed, AID-tagged DAF-16 was not detectable in TIR1-expressing neurons (all pharyngeal neurons except I1 and I4) after auxin treatment. Scale bars: 20 μm. (B) Pharyngeal pumping rate after DAF-16/FoxO depletion in *daf-2(e1370)* dauers using *ehs-1prom4*-driven TIR1 (*otEx7628* and *otEx7629*). Horizontal lines represent median for 36 animals per condition. *** indicates *p* < 0.001 in Mann-Whitney test. (C) DAF-16/FoxO depletion in glutamatergic pharyngeal neurons of *daf-2(e1370)* dauers using *eat-4prom7*-driven TIR1. In 20/20 animals observed, AID-tagged DAF-16 was not detectable in TIR1-expressing pharyngeal neurons (I5 and M3) after auxin treatment. Scale bars: 20 μm. (D) Pharyngeal pumping rate after DAF-16/FoxO depletion in *daf-2(e1370)* dauers using *eat-4prom7*-driven TIR1 (*otEx7670* and *otEx7671*). Horizontal lines represent median for 36 animals per condition. **** indicates *p* < 0.0001 in Mann-Whitney test. (E) DAF-16/FoxO depletion in cholinergic pharyngeal neurons of *daf-2(e1370)* dauers using *unc-17prom4*-driven TIR1. In 20/20 animals observed, AID-tagged DAF-16 was not detectable in TIR1-expressing pharyngeal neurons (I1, M1, M2, M4, M5 and MC) after auxin treatment. Scale bars: 20 μm. (F) Pharyngeal pumping rate after DAF-16/FoxO depletion in *daf-2(e1370)* dauers using *unc-17prom4*-driven TIR1 (*otEx7668* and *otEx7669*). Horizontal lines represent median for 36 animals per condition. ** indicates *p* < 0.01 in Mann-Whitney test. (G) Neurotransmitter identity of all 14 pharyngeal neurons. Shaded boxes for each promoter fragment indicate the neurons in which TIR1 expression is strong enough to completely deplete AID-tagged DAF-16/FoxO in the presence of auxin (20 animals observed per strain). (**H** and **I**) Pharyngeal pumping rate in *daf-2(e1370)* dauers transferred from ethanol (solvent) to auxin or from auxin to ethanol after dauer entry (dauers were transferred on day 3 after hatching, pumping rate was measured on day 6 after hatching). DAF-16/FoxO was depleted in the intestine (*Pges1::TIR1*) or in no tissue (no TIR1 control) in the presence of auxin. Horizontal red lines represent median for 18 animals per condition. *p* values are for Dunn’s multiple comparisons test performed after one-way ANOVA on ranks.

We conclude that DAF-16/FoxO function is required in two distinct tissues to control behavioral and structural remodeling of the pharynx, the nervous system as well as pharyngeal muscle. As detailed in the next section, we found yet another tissue type that requires DAF-16/FoxO for pharynx remodeling.

### Intestinal removal of DAF-16/FoxO has cell-autonomous and cell non-autonomous consequences

We further explored autonomous and non-autonomous functions of DAF-16/FoxO by depleting it from the intestine using a *ges-1-*driven TIR1 transgene. In a *daf-2* mutant background, these transgenic animals still enter the dauer stage, indicating that DAF-16/FoxO is not required in the intestine for dauer entry (**Fig.4B**). The lack of an effect on dauer formation allowed us to study the function of DAF-16/FoxO in the intestine of dauer stage animals.

Using Oil Red O staining, we observed that DAF-16/FoxO is required in the intestine to control the metabolic remodeling of intestinal cells observed upon entry into the dauer stage (**Fig.10A,B**). This is consistent with previous studies that utilized transgenic rescue approaches (Zhang et al., 2013) and with the fact that the promoter of intestinally expressed fat metabolism genes are directly targeted by DAF-16/FoxO (Murphy et al., 2003; Oh et al., 2006).

**Fig.10.**
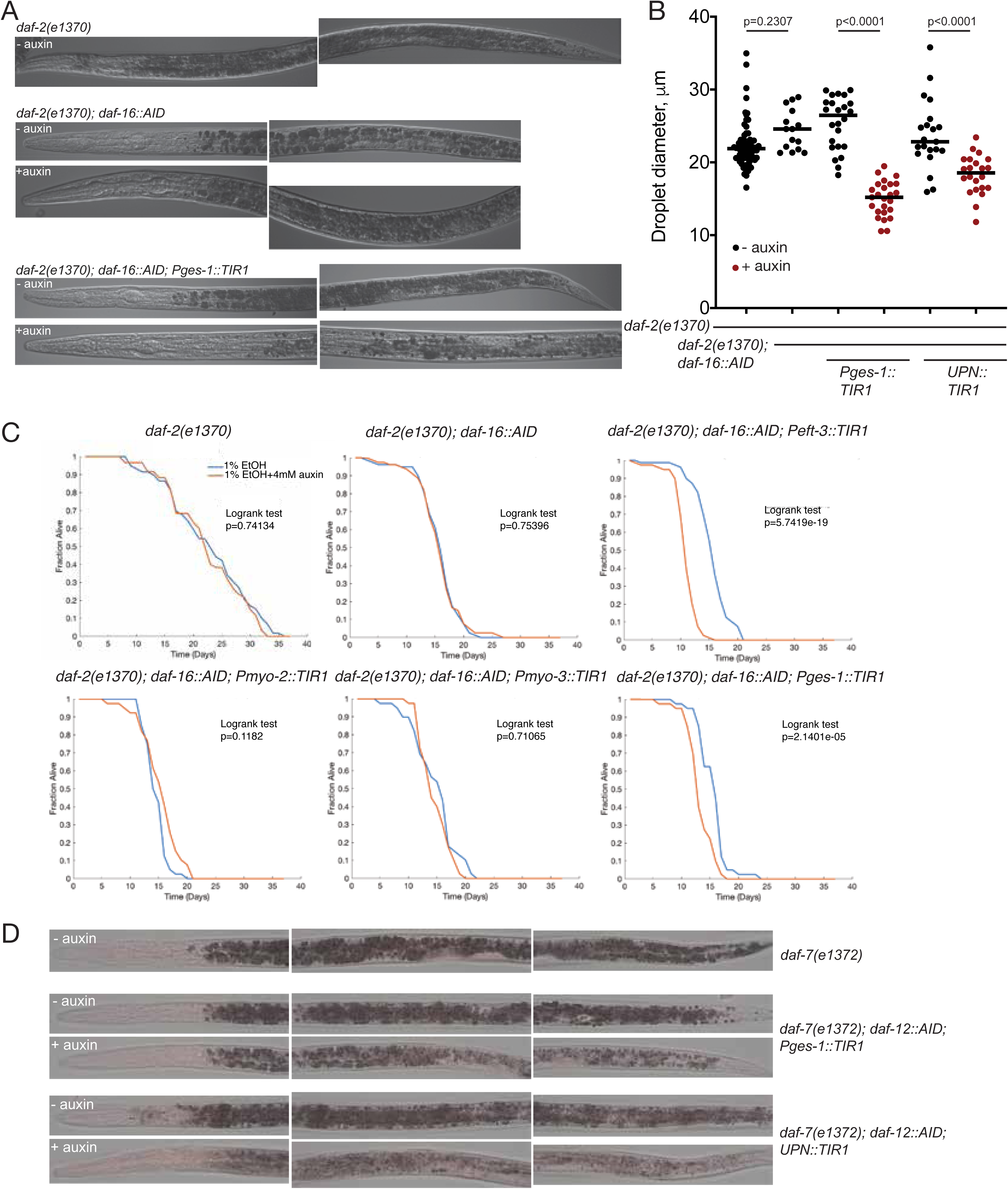
**Intestinal removal of DAF-16/FoxO and DAF-12 has cell autonomous and cell non-autonomous consequences** (A) Oil Red O staining of control and intestinal DAF-16/FoxO-depleted strains. Note a decrease in the intensity of staining upon auxin treatment of the strain with intestinal depletion of DAF-16/FoxO. (B) Quantification of lipid droplet size for strains with intestinal and panneuronal depletion of DAF-16/FoxO reveals autonomous and non-autonomous regulation of dauer lipid metabolism. (C) Survival curves for control and auxin treatment conditions for the strains of the indicated genotypes. (**C1-2**) Long lifespan of *daf-2(e1370)* mutants requires DAF-16. (**C3**) Ubiquitous depletion of DAF-16/FoxO strongly reduces lifespan in a *daf-2(e1370)* background. (**C4-5**) Depletion of DAF-16/FoxO in pharyngeal or body wall muscles did not significantly reduce lifespan in a *daf-2(e1370)* background. (**C6**): Intestinal depletion of DAF-16/FoxO strongly reduces lifespan in a *daf-2(e1370)* background. N = 40-80 worms for each group. (D) Oil Red O staining of control and intestinal and panneuronal DAF-12/VDR-depleted strains. Panneuronal DAF-12/VDR depletion has a stronger effect on dauer lipid reserves.

Loss of function mutants for the insulin/IGF receptor ortholog DAF-2 exhibit longer lifespan than wild type animals, an effect that depends on DAF-16/FoxO (Kenyon et al., 1993). We recapitulated this effect through DAF-16::AID removal with a ubiquitously expressed TIR1 driver (**Fig.10C**). Complementing previous transgenic rescue studies (Libina et al., 2003), we find that depletion of DAF-16/FoxO exclusively from the intestine strongly suppresses the lifespan extension of *daf-2(e1370)* mutants (**Fig.10C**), while muscle--specific *daf-16* removal does not affect life span (**Fig.10C**). We noted that DAF-16::mNG::AID animals already have a reduced lifespan on their own even in the absence of TIR1 but nevertheless, in the presence of ubiquitous or intestinal TIR1 and auxin its effects on *daf-2* longevity become strongly enhanced (**Fig.10C**).

Unexpectedly, we discovered that in addition to its metabolic and lifespan effect, intestinal DAF-16/FoxO depletion also results in a complete failure to remodel pharyngeal pumping behavior and pharyngeal muscle constriction (**Fig.8A, B**). By auxin-shifting the dauers initially grown in control conditions, we find that the DAF-16/FoxO requirement in the gut is continuous, since removal of DAF-16/FoxO in the intestine post-dauer entry still results in derepression of pharyngeal pumping (**Fig.9H, I**). Taken together, DAF-16/FoxO is required in three different tissue types to control pharyngeal remodeling, one of them (the intestine) being completely external to the pharynx itself.

We also asked whether intestinal DAF-16/FoxO has other non-autonomous defects in the nervous system. We specifically asked whether the downregulation of *inx-2*, which we found not to required neuronal DAF-16/FoxO (see above; **Fig.6E**), may require intestinal DAF-16/FoxO. We found this not to be the case (**Suppl. Fig.S5**). Similarly, intestinal DAF-16/FoxO depletion does not affect the neuronal changes of *che-7* and *inx-6* induction in neuronal cells (**Suppl. Fig.S5**).

We conclude that intestinal depletion of DAF-16/FoxO has both autonomous effects in the intestine, but also non-autonomous effects in non-intestinal tissue. We also found that the intestine is at the receiving end of DAF-16/FoxO*-*mediated non-cell autonomous function in other tissues. Specifically, panneuronal depletion of DAF-16/FoxO affects the size of Oil Red O-positive droplets in the intestine (**Fig.10B**). In these animals, intestinal DAF-16/FoxO still translocated normally to the nucleus (**Suppl. Fig.S1**), indicating that (a) neuronal DAF-16/FoxO is not providing insulin-mediated signals to promote nuclear translocation of intestinal DAF-16/FoxO, and (b) that intestinal nuclear translocation of DAF-16/FoxO is alone not sufficient to affect Oil Red O droplets size in the intestine.

### DAF-16/FoxO acts cell-autonomously for muscle remodeling

Muscles also undergo remodeling upon entry in the dauer stage. They extend additional muscle arms into the nerve cords to receive supernumerary synaptic inputs from motor neurons, leading to an increased sensitivity to neurotransmitter signaling (Dixon et al., 2008; Lewis et al., 1987). Using a muscle-specific TIR1 driver line, we find that DAF-16/FoxO is required in muscle during dauer remodeling to generate these supernumerary muscle arms (**Fig.11A, B**).

**Fig.11.**
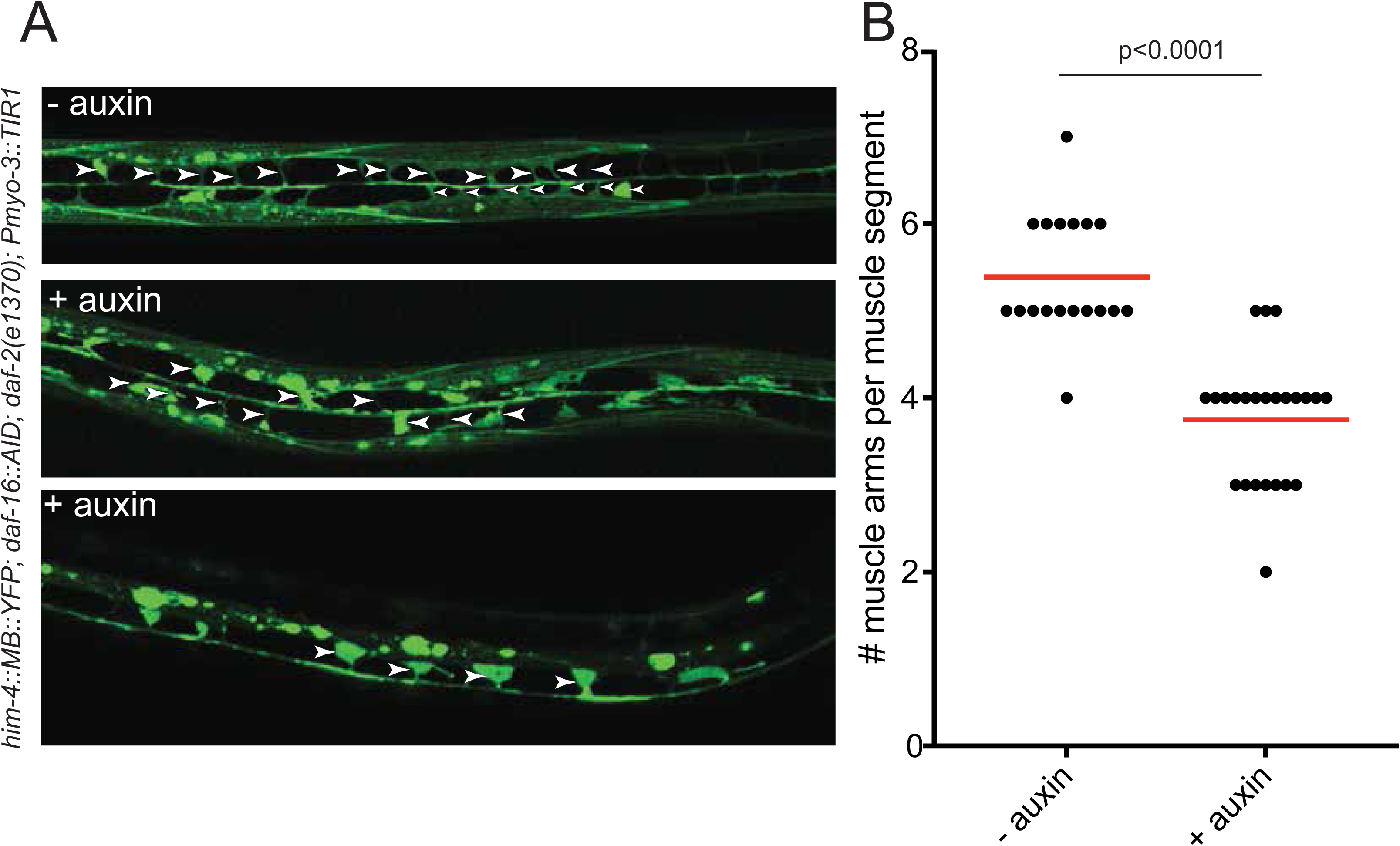
**DAF-16/FoxO acts cell-autonomously in muscle remodeling** (A) Depleting DAF-16/FoxO from body wall muscle results in fewer muscle arms than in control dauers, as visualized with the *trIs30[him-4p::MB::YFP]* reporter (Dixon and Roy, 2005). (B) Quantification of the number of muscle arms per muscle segment in control and auxin-treated dauers.

Body wall muscle of dauer animals dramatically shrink, as determined by electron microscopy (https://www.wormatlas.org/dauer/muscle/Musframeset.html). We find that muscle-specific depletion of DAF-16/FoxO affected overall dauer morphology, resulting in a widening of the body and a slight increase in body length (**Suppl.Fig.6A-C**). This observation is consistent with DAF-16/FoxO acting cell autonomously in muscle to affect muscle shrinkage and hence, overall body width. As mentioned above, muscle-specific removal of DAF-16/FoxO has no effect on locomotion of the animals (**Suppl. Fig.S6**).

### Insulin-FoxO signaling is sufficient to suppress pharynx pumping

The above DAF-16/FoxO depletion experiments demonstrate that DAF-16/FoxO is required in a number of distinct tissue types to suppress tissue remodeling. We asked whether insulin signaling-controlled DAF-16/FoxO may also be sufficient to control remodeling. To this end, we generated a dominant negative version of the insulin/IGF-like receptor DAF-2 (“DAF-2^DN^”) by replacing its intracellular kinase domain with a fluorophore (**Fig.12A**). Due to dimerization properties of insulin/IGF-like receptors, DAF-2^DN^ is predicted to antagonize endogenous receptor signaling through the formation of an inactive dimer; this should then result in a nuclear translocation of DAF-16/FoxO. We overexpressed this DAF-2^DN^ construct in three different tissue types, in which we had found DAF-16/FoxO to be required to downregulate pharyngeal pumping (panneuronal, intestinal and pharyngeal muscle). We find that in young adult animals, in which we express DAF-2^DN^ panneuronally, DAF-16/FoxO partially translocates to neuronal nuclei, as expected (**Fig.12B**). These animals display a significantly reduced pharyngeal pumping rate (**Fig.12C**), demonstrating that the inhibition of insulin signaling and hence activation of DAF-16/FoxO is sufficient to downregulate pumping, even outside of dauer context. The same phenotypic consequences are observed upon intestinal expression of DAF-2^DN^. In these animals, DAF-16/FoxO translocates to the nucleus in the intestine, and there is a concomitant downregulation of pharyngeal pumping (**Fig.12B, C**). Unexpectedly, pharyngeal muscle expression of DAF-2^DN^ does not trigger nuclear translocation in pharyngeal muscle (**Fig.12B**). Consistent with this, pharyngeal muscle-expressed DAF-2^DN^ does not result in a reduction of pharyngeal pumping (**Fig.12C**).

**Fig.12.**
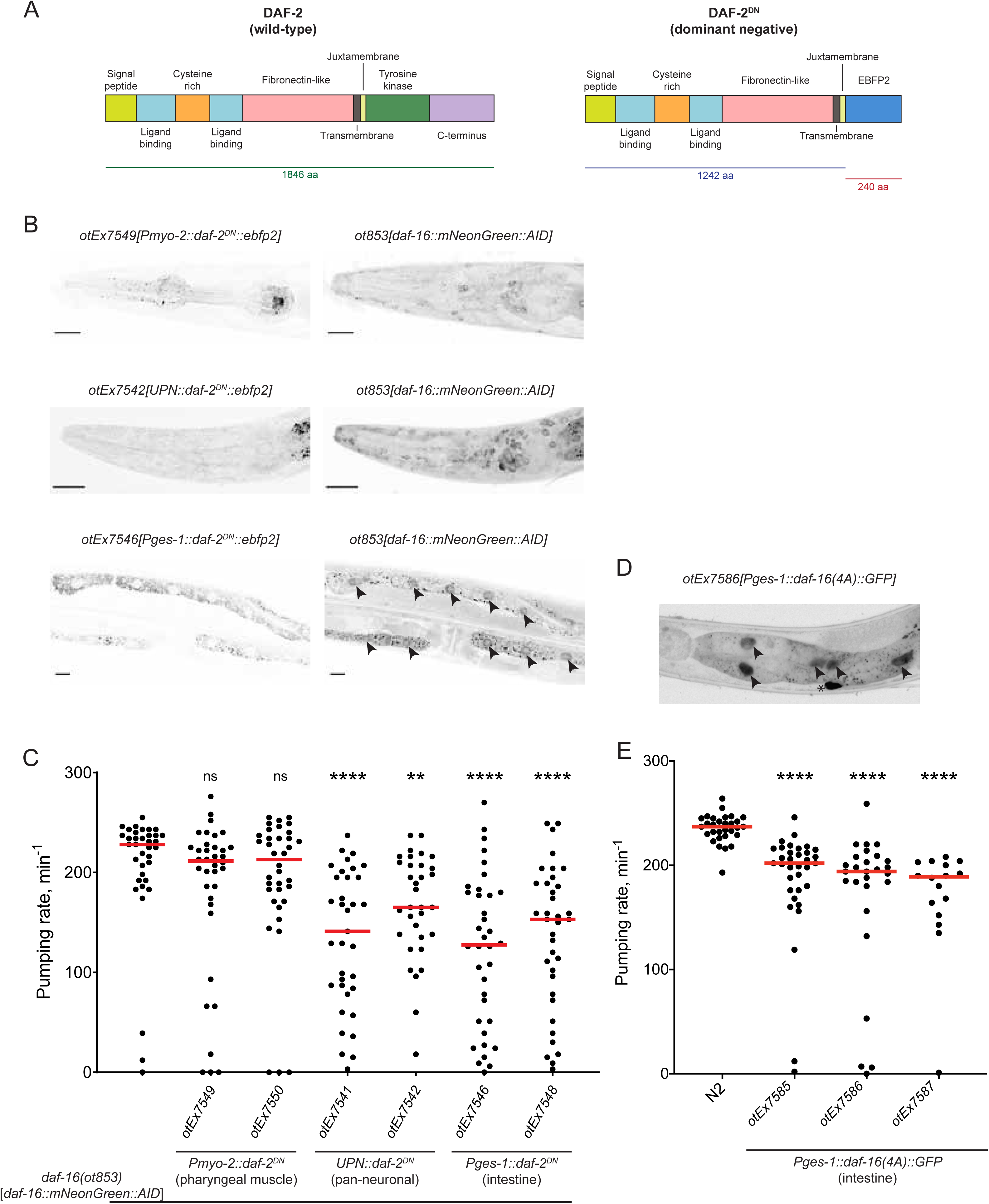
**DAF-16/FoxO activation via inhibition of insulin signaling is sufficient to cell non-autonomously reduce pharyngeal pumping** (A) Schematic showing protein domains of wild-type DAF-2 (*left*) and DAF-2^DN^, a dominant negative form of DAF-2/InsR in which the tyrosine kinase and C-terminus domains are replaced with the blue fluorescent protein EBFP2 (*right*). (B) *Left panels*: Expression of *daf-2^DN^::ebfp2* in pharyngeal muscle (*top*), pan-neuronal (*middle*) and intestinal (*bottom*) tissues. *Right panels*: Localization of endogenously mNeonGreen-tagged DAF-16 protein in the same animals shown in the corresponding panels on the left. Black arrowheads show nuclear localization of DAF-16 in intestinal tissue. Scale bars: 20 μm. (C) Pharyngeal pumping rate in day 1 adult animals expressing *daf-2^DN^::ebfp2* in pharyngeal muscle (*otEx7549* and *otEx7550*), pan-neuronal (*otEx7541* and *otEx7542*) and intestinal (*otEx7546* and *otEx7548*) tissues in the endogenously mNeonGreen-tagged *daf-16(ot853)* genetic background. Red horizontal lines represent median for ≥ 35 animals per condition. **, **** and ns indicate *p* < 0.01, *p* < 0.0001 and *p* ≥ 0.05 compared to the *daf-16(ot853)* control strain in Dunn’s multiple comparisons test performed after one-way ANOVA on ranks. (D) Expression of GFP-tagged DAF-16(4A) in intestinal tissue of animals. Black arrowheads show strong nuclear localization of DAF-16(4A)::GFP. Asterisk indicates non-specific autofluorescence signal. (E) Pharyngeal pumping rate in day 1 adult wild-type (N2) strain and transgenic animals expressing *daf-16(4A)::gfp* in intestinal tissue (*otEx7585-87*). Red horizontal lines represent median for ≥ 16 animals per condition. **** indicates *p* < 0.0001 compared to wild-type strain in Dunn’s multiple comparisons test performed after one-way ANOVA on ranks.

As a complementary approach to the DAF-2^DN^ experiments, we also generated transgenic animals that express an activated, insulin-signaling-independent form of DAF-16/FoxO exclusively in the intestine. This activation was achieved by mutating four residues (serine or threonine) that are normally phosphorylated by Akt kinases to prevent DAF-16/FoxO nuclear translocation in fed animals (Lin et al., 2001; Paradis and Ruvkun, 1998). We confirmed that in these animals, mutated DAF-16/FoxO is indeed now localized to the nucleus (**Fig.12D**), and we find that in normal, non-starved adult animals, pharyngeal pumping is significantly reduced (**Fig.12E**).

### The nuclear hormone receptor DAF-12/VDR also controls neuronal remodeling

Even though the nuclear hormone receptor *daf-12/VDR* has previously been shown to be essential for dauer remodeling (Antebi et al., 1998; Antebi et al., 2000), there have been no studies that address its focus of action for the dauer decision and/or dauer-specific tissue remodeling events. We first asked whether and where DAF-12/VDR is required for the remodeling of locomotory behavior in dauer animals. We used the same strategy as for DAF-16/FoxO and found that panneuronal depletion of DAF-12/VDR results in striking deviations from control dauer locomotion, even stronger than those observed upon panneuronal depletion of DAF-16/FoxO (**Fig.7C, D**). Specifically, panneuronally DAF-12/VDR-depleted dauers crawl faster, pause less, coil and turn more frequently, and have a flatter waveform (decreased wave amplitude) than control dauers (**Fig.7C, D**, **Suppl.Fig.6A, B**). All these features make them more L3-like, as a comparison with L3 larvae of the background *daf-7(e1372)* strain shows. The magnitude of effect in many cases is larger than that of DAF-16/FoxO panneuronal depletion (**Fig.7B, D**).

Panneuronal DAF-12/VDR depletion also affects pharyngeal pumping and alterations in GPCR expression profiles. As with the locomotory effect, DAF-12/VDR shows some, but not complete overlaps in cellular specificity with DAF-16/FoxO. Dauer-specific expression changes of *sri-9* require DAF-16/FoxO and DAF-12/VDR in the NSM, OLL, AIZ, AWC neurons (**Fig.5A, D**). However, DAF-12/VDR is not required for *sri-9* changes in the ADA and AIM neurons (**Fig.5D**).

There are also cell non-autonomous functions of neuronal DAF-12/VDR. We find that DAF-12/VDR removal from the nervous system affects pharyngeal muscle constriction and intestinal metabolic remodeling (**Fig.8D, E; Fig.10D**). However, unlike the case of DAF-16/FoxO, pharyngeal muscle depletion of DAF-12/VDR does not affect pharyngeal pumping or pharyngeal muscle constriction (**Fig.8D, E**).

As in the case with DAF-16/FoxO, intestinal DAF-12/VDR depletion does not prevent dauer formation (in the *daf-7* mutant background) (**Fig.4C**). In contrast to the intestinal DAF-16/FoxO depletion, intestinal DAF-12/VDR depletion does not affect pharyngeal muscle constriction or pharyngeal pumping. However, intestinal DAF-12/VDR depletion does affect Oil red O straining in the intestine, but not as strongly as panneuronal DAF-12/VDR depletion does (**Fig.10B**)

Taken together, similar to DAF-16/FoxO, DAF-12/VDR displays cell-autonomous functions in a number of neuronal remodeling events, but also has non-autonomous roles, with the phenotypic spectrum of the tissue-specific TF depletion overlapping in some, but not in other, cases.

### The cellular specificity of the effect of the broadly expressed hormonal systems is controlled by terminal selector-type transcription factors

DAF-16/FoxO and DAF-12/VDR are both broadly expressed but the read-outs of their activity are highly cell-type specific. For example, GPCR- or innexin-encoding genes are turned on in a DAF-16-dependent manner in only specific sets of neurons. To address the cell-type and target specificity of the effects of DAF-16/FoxO and DAF-12/VDR, we considered a potential collaboration with terminal selector-type TFs, master regulatory TFs that initiate and maintain neuron-type specific gene expression programs during nervous system development (Hobert, 2016). For example, DAF--16/FoxO-dependent upregulation of a fosmid-based *che-7* reporter in the BDU, NSM or AIM neurons may be the result of *daf-16* cooperating with the terminal selector of BDU, NSM and AIM neuron identity, the *unc-86* POU homeobox gene (which acts in distinct cofactor combinations in these different neuron types). To test this hypothesis, we asked whether the dauer-specific upregulation of *che-7* in BDU, NSM and AIM requires *unc-86*. Using *unc-86* null mutant animals, we indeed found this to be the case (**Fig.13A, B**). Similarly, the DAF- 16/FoxO*-*dependent upregulation of the GPCR *sri-9* in the NSM neuron also requires *unc-86* (**Fig.13C**). In conclusion, the cellular specificity of DAF-16/FoxO activity appears to be dictated by the neuron-type specific complement of terminal selectors.

**Fig.13.**
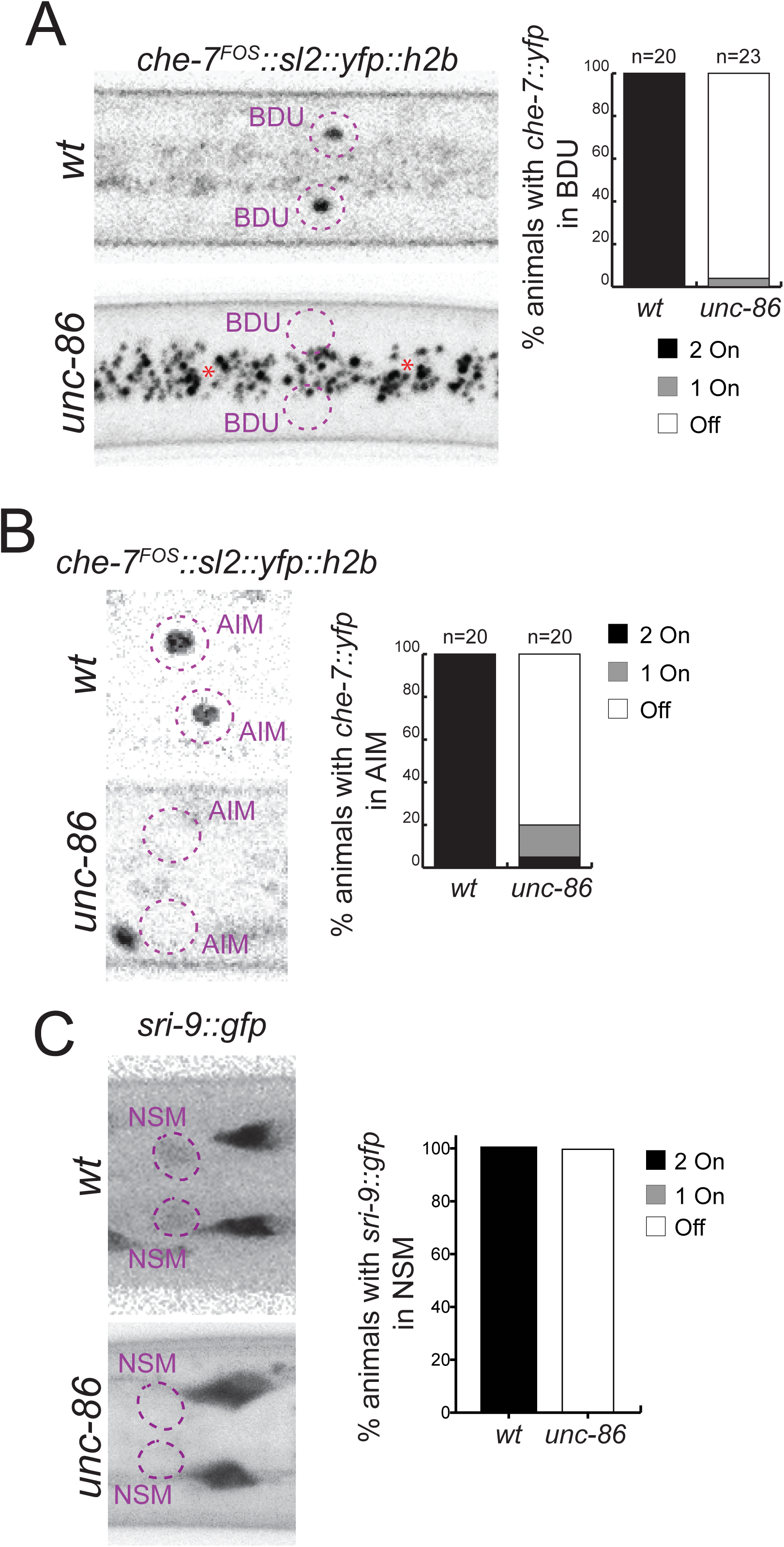
**Dauer-specific expression of neuronal reporters depends on terminal selector-type transcription factors** (A) Dauer-specific expression of a *che-7* fosmid reporter (*otEx7112*) in BDU neurons is abolished in *unc-86(n846)* mutants. (B) Dauer-specific expression of a *che-7* fosmid reporter (*otEx7112*) in AIM neurons is abolished in *unc-86(n846)* mutants. (C) Dauer-specific expression of a *sri-9* reporter (*otIs732*) in NSM neurons is abolished in *unc-86(n846)* mutants.

## DISCUSSION

The ability of cell types to undergo a change in phenotype (“cellular plasticity”) is a hallmark of many different cell types throughout the animal kingdom. How such cellular plasticity is controlled on a single cell level and how such plasticity is coordinated through many tissue types of an organism are fascinating questions that we address here in this paper. The *C. elegans* dauer-remodeling paradigm highlights the importance of hormonal signaling events. Classic genetic epistasis analysis has revealed complex relationships between these hormonal signaling events, but many previous studies have focused on relatively crude binary read-outs (whether animals go into dauer or not) and/or used experimental approaches that entailed certain limitations (most notably, problematic overexpression approaches that could have neomorphic effects; (Bansal et al., 2014)). We have used the CRISPR genome engineering technology to generate expression reagents and conditional alleles that allowed us to revisit a number of questions about the manner in which this hormonal control defines cellular and behavioral remodeling events. Our genetic loss-of-function approaches have probed the genetic requirement of the key effector TFs of these hormonal signaling systems and are orthogonal to previous rescue approaches, which probed the sufficiency, rather than necessity, of these hormonal effector systems. Previous genetic mosaic analysis, particularly of the TGFβ (Inoue and Thomas, 2000) and insulin-signaling systems (Apfeld and Kenyon, 1998), in the context of dauer formation, are conceptually similar to our tissue-specific protein depletion approach, but the latter has provided greater control over the cell types in which a specific protein is depleted, as well as about the timing of the action of these systems. We will first discuss timing and then discuss the focus of action.

The temporal control of gene activity achieved through the AID system allowed us to address a previously unresolved question – are these hormonally controlled TF effectors of dauer remodeling only transiently required for entry into the dauer stage or is their activity continuously required? Since the vertebrate FoxO proteins have been proposed to act as pioneer transcription factors (Zaret and Carroll, 2011), a transient activity of at least DAF-16/FoxO appeared plausible, but our explicit demonstration of a continuous requirement of DAF-16/FoxO, as well as of DAF-3/SMAD and DAF-12/VDR, rather suggest that these TFs are likely continuously engaged on their target gene promoters to actively maintain the dauer state.

In regard to the focus of action of the three hormonal systems, our analysis (summarized in **Fig.14**) has confirmed two key previously made conclusions about the focus of action of the TGFβ and insulin-signaling systems: (1) TGFβ signaling via its effector TF DAF-3/SMAD operates within the nervous system to non-autonomously control tissue remodeling throughout the entire animal. We have extended this observation by demonstrating that TGFβ signaling is specifically required within sensory neurons, perhaps for the expression of insulin ligands that then signal to other cell types. (2) Insulin-signaling via its effector TF DAF-16/FoxO operates cell autonomously in a variety of distinct target tissues to control cellular remodeling. We have explicitly demonstrated such autonomy of DAF-16/FoxO function in a number of different tissue types, most extensively in the nervous system, where DAF-16/FoxO controls a plethora of instances of neuronal phenotypic plasticity. Our analysis of DAF-16/FoxO*-*depleted neurons was enabled by the unanticipated observation that neuronal DAF-16/FoxO depletion does not affect entry into the dauer stage. This observation was unanticipated because previous work has shown that resupplying *daf-16* in the nervous system rescues the Daf-c-suppression phenotype of *daf-16* mutants (Libina et al., 2003). However, we cannot exclude the possibility that the AID system may not remove all protein function. Arguing against such possibility is that (a) in all cases examined, we do observe efficient tagged fluorescent protein removal; (b) AID-mediated DAF-16/FoxO depletion does phenocopy the null phenotype when ubiquitously implemented and (c) AID-mediated neuronal DAF-16/FoxO depletion does result in a number of neuronal remodeling defects, indicating successful removal of DAF-16/FoxO. In conclusion, while the site of function for DAF-16/FoxO control of the entry into the dauer stage remains unresolved, our analysis clearly shows that DAF-16/FoxO operates cell-autonomously in the nervous system to control neuronal remodeling.

**Fig.14.**
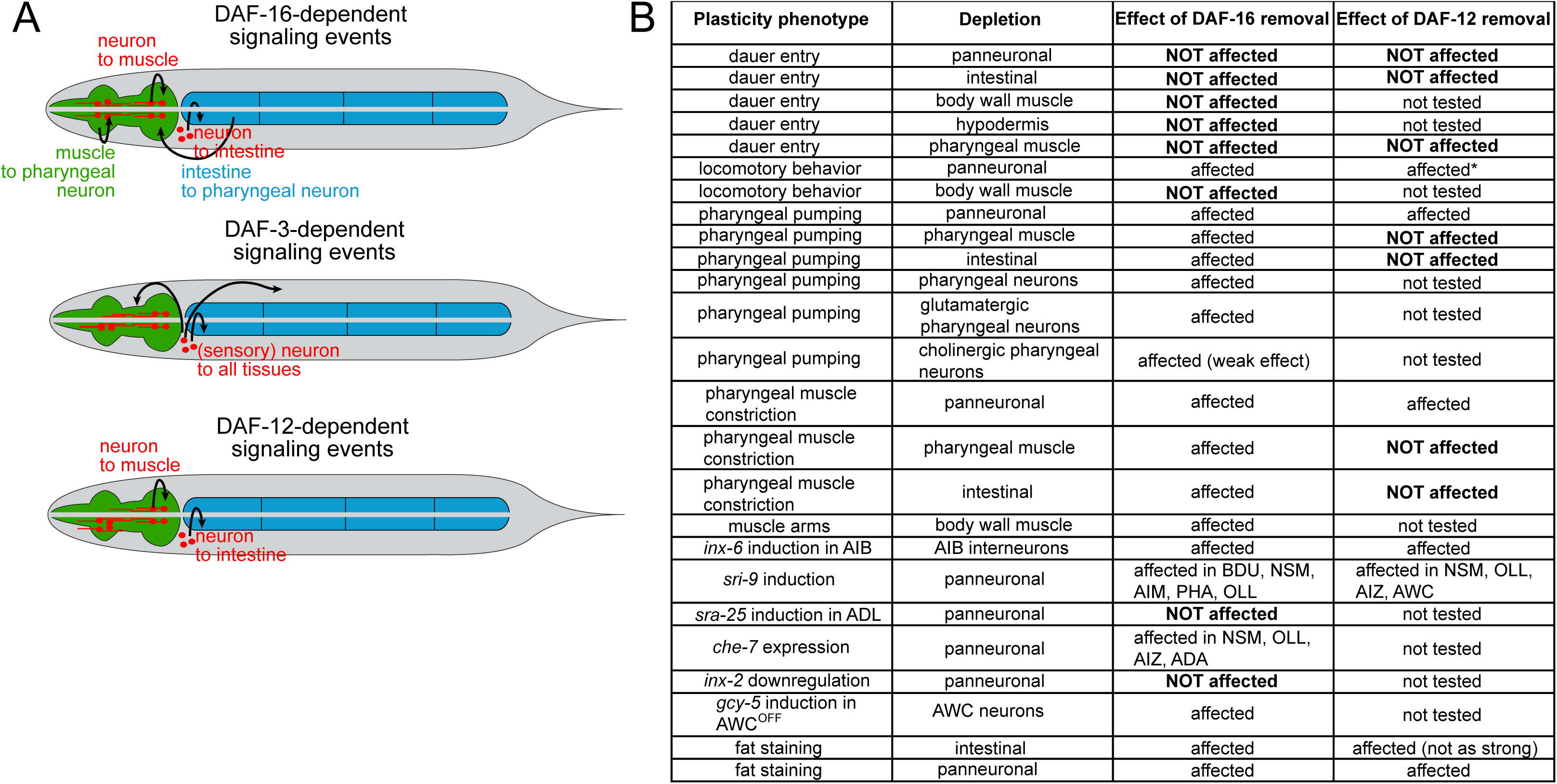
**Model and summary** (A) Models of inter-tissue signaling in dauer formation for DAF-16/FoxO, DAF-3/SMAD and DAF-12/VDR (B) Table summarizing differences in phenotypic outcomes of tissue-specific depletion of DAF-16/FoxO and DAF-12/VDR

In addition, our analysis revealed a plethora of unexpected cell non-autonomous activities of DAF-16/FoxO and DAF-12/VDR that indicates the existence of several interorgan signaling axes (summarized in **Fig.14A, B**). While neuronal DAF-16/FoxO may display a cell-autonomous requirement for pharyngeal pumping behavior, neuronal DAF-16/FoxO also affects the remodeling of pharyngeal muscle, as well as the metabolic remodeling of intestinal tissue observed upon dauer entry. Neuropeptidergic signaling axes from the nervous system to the gut have been described (Lemieux and Ashrafi, 2015; Srinivasan, 2020), and it is possible that during the dauer stage, these axes become modulated in the nervous system by DAF-16/FoxO. How neuronal DAF-16/FoxO affects the autophagy-dependent constriction of pharyngeal musculature is presently entirely unclear, as is the manner by which intestinal DAF-16/FoxO affects pharyngeal muscle constriction and behavior. Notably, pharyngeal muscle constriction and cessation of pumping behavior are not obligatorily linked. Constricted pharynxes can be made to pump (**Suppl.Fig.7C**) and, vice versa, pharyngeal pumping can cease without muscle constriction (Keane and Avery, 2003).

Another intriguing interorgan signaling axis that we described here is from the intestine to the pharyngeal nervous system, resulting in a downregulation of pharyngeal pumping (**Fig.14A**). Our experiments indicate that DAF-16/FoxO is not only required, but also sufficient to instruct the intestine to control the activity of the pharyngeal nervous system. Since DAF-16/FoxO also acts within the pharyngeal nervous system, we hypothesize that intestinal DAF-16/FoxO modulates the release of an insulin signal from the intestine that controls DAF-16/FoxO activity in the pharyngeal neurons to control pumping behavior. Similarly, the effect of neuronal DAF-16/FoxO on intestinal DAF-16/FoxO-dependent metabolic remodeling suggests the DAF-16/FoxO-dependent release of insulin from the nervous system. The abundance and complex expression patterns of insulin-encoding genes in *C. elegans* (Duret et al., 1998; Ritter et al., 2013) provide plenty of room for such multidirectional “insulin relay” systems (**Fig.14A**).

Lastly, our tissue-specific DAF-12/VDR removal experiments provide insights into the focus of DAF-12/VDR action in tissue remodeling and its relationship to DAF--16/FoxO. The genetic relationship of these two TF systems is complex and has mostly been studied in the context of the developmental arrest or organismal ageing (Gottlieb and Ruvkun, 1994; Jeong et al., 2010; Matyash et al., 2004; Rottiers et al., 2006; Vowels and Thomas, 1992). We have shown here a range of striking similarities in the effect of cell-type specific depletion of DAF-16/FoxO or DAF-12/VDR on the remodeling of distinct tissue types. Both transcription factors act autonomously in the nervous system for neuronal remodeling events and non-autonomously in the nervous system to affect metabolic remodeling of the intestine. It is conceivable that they interact on the level of a target gene promoter to, for example, promote the expression of an insulin gene that may affect intestinal remodeling. However, intestinal depletion of DAF-12/VDR, while also affecting intestinal remodeling, does not affect pharyngeal remodeling. This indicates a distinct target gene spectrum of DAF-16/FoxO and DAF-12/VDR in the intestine, with only DAF-16/FoxO controlling the expression of a signal to the pharyngeal nervous system.

The mode of interaction between DAF-12/VDR and DAF-16/FoxO in the nervous system is presently unclear. While there are many phenotypic similarities between DAF-12/VDR and DAF-16/FoxO, the phenotypes of DAF-12/VDR generally appear more restricted. Even the effects on the same target gene shows cell-type specific differences in DAF-12/VDR and DAF-16/FoxO-dependence. The upregulation of the GPCR *sri-9* requires both DAF-12/VDR and DAF-16/FoxO in four neuron classes, but only DAF-16/FoxO in an additional two neuron classes. The different spectra of activity argue that DAF-16/FoxO and DAF-12/VDR do not obligatorily act together, but these two TFs can have their own independent functions.

It is fascinating to consider how these plasticity phenomena intersect with the genetically hardwired differentiation programs of individual neuron programs, executed right after the birth of a neuron. These hardwired differentiation programs are controlled by terminal selector-type TFs (Hobert, 2016). Generally, one emerging theme is that both hormonal systems intersect with the ability of terminal selectors to exert their functions. For example, the dauer-specific upregulation of a number of effector genes, such as the *inx-6* gene in AIB or the *che-7* gene in BDU, require a cell-type specific terminal selector transcription factor together with the DAF-16/FoxO (UNC-42/Prop1 in AIB or UNC-86/Brn3 in BDU)(Bhattacharya et al., 2019)(this paper). In the case of *inx-6* and AIB, we find that not only *daf-16* but also *daf-12* is required for the *unc-42-*dependent upregulation of *inx-6.* It is presently not clear whether this is an indication of DAF-12/VDR operating as a transcriptional activator or whether it operates indirectly via a repressive mechanism.

All the signaling systems that we described here to affect cellular plasticity in the nervous system and beyond have vertebrate counterparts that have also been shown to affect brain physiology and plasticity. The vertebrate DAF-12 homolog VDR is, like DAF-12, broadly expressed in the brain and mediates the effects that vitamin D has on neuronal development and plasticity (Cui et al., 2017). Similarly, insulin and insulin-like peptides have been shown to affect neuronal plasticity in a variety of different contexts (Fernandez and Torres-Aleman, 2012; Ferrario and Reagan, 2018). In any of these systems, including *C. elegans*, it will be fascinating to explore through what effector genes these hormonal signals operate and how the cellular-specificity of such effector systems is controlled.

## MATERIAL AND METHODS

### Mutant and transgenic strains

The following mutant strains of *C. elegans* were used in this study:

CB1370 *daf-2(e1370)*

CB1372 *daf-7(e1372)*

MT1859 *unc-86(n846)*

CB1338 *mec-3(e1338)*

DR26 *daf-16(m26)*

The following transgenic strains of *C. elegans* were used in this study:

BC13401 *sIs12199 [sra-25::GFP]* (Hunt-Newbury et al., 2007)

OH3333 *otIs178[flp-4::gfp]*

OH13405 *ot804 [inx-6::sl2::yfp::H2B]* (Bhattacharya et al., 2019)

OH13102 *otIs586[gcy-5::NLS::GFP]* (Patel and Hobert, 2017)

OH13908 *ot821[daf-16::mKate2::3xFlag]*(Aghayeva et al., 2020)

OH14125 *ot853 [daf-16::mNG::3xFlag::AID]* (Bhattacharya et al., 2019)

OH14654 *ot853 [daf-16::mNG::3xFlag::AID]; daf-2(e1370)*

OH14888 *ot853 [daf-16::mNG::3xFlag::AID]; daf-2(e1370); ieSi57 [Peft-3::TIR1::mRuby]*

OH14897 *ot853 [daf-16::mNG::3xFlag::AID]; daf-2(e1370); ieSi60 [Pmyo-2::TIR1::mRuby]*

OH14898 ot853*[daf-16::mNG::3xFlag::AID]; daf-2(e1370); otTi1 [Pmyo-3::TIR1::mRuby]*

OH14945 *ot853 [daf-16::mNG::3xFlag::AID]; daf-2(e1370); ieSi61 [Pges-1::TIR1::mRuby]*

OH14915 *ot853 [daf-16::mNG::3xFlag::AID]; daf-2(e1370); otTi27 [Pdpy-7::TIR1::mTur2]*

OH15845 *ot853 [daf-16::mNG::3xFlag::AID]; daf-2(e1370); otIs730 [Punc-11prom8+ehs-1prom7+ rgef1prom2::TIR1::mTur2]*

OH16029 *ot975[daf-16::mNeptune2.5::3xFlag::AID]* (Aghayeva et al., 2020)

OH14892 *ot875[daf-3::GFP::3xFlag]*

OH14896 *ot877 [daf-3::TagRFP::3xFlag AID]*

OH14985 *ot877 [daf-3::TagRFP::3xFlag AID]; daf-7(e1372)*

OH14946 *ot877 [daf-3::TagRFP::3xFlag::AID]; daf-7(e1372); ieSi57 [Peft-3::TIR1::mRuby]*

OH14948 *ot877 [daf-3::TagRFP::3xFlag::AID]; daf-7(e1372); ieSi60 [Pmyo-2::TIR1::mRuby]*

OH15914 *ot877 [daf-3::TagRFP::3xFlag::AID]; daf-7(e1372); otIs730 [Punc-11prom8+ehs-1prom7+ rgef1prom2::TIR1::mTur2]*

OH14589 *ot870 [daf-12::GFP::3xFlag]*

OH14891 *ot874 [daf-12::TagRFP::3xFlag::AID]*

OH14984 *daf-7(e1372); ot874 [daf-12::TagRFP::3xFlag::AID]*

OH14986 *ot874 [daf-12::TagRFP::3xFlag::AID]; daf-7(e1372); ieSi57 [Peft-3::TIR1::mRuby]*

OH14988 *ot874 [daf-12::TagRFP::3xFlag::AID]; daf-7(e1372); ieSi61 [Pges-1::TIR1::mRuby]*

OH14989 *ot874 [daf-12::TagRFP::3xFlag::AID]; daf-7(e1372); ieSi60 [Pmyo-2::TIR1::mRuby]*

OH15913 *ot874 [daf-12::TagRFP::3xFlag::AID]; daf-7(e1372); otIs730 [Punc-11prom8+ehs-1prom7+ rgef1prom2::TIR1::mTur2]*

OH15597 *ot804[inx-6::SL2::YFP::H2B]; ot853 [daf-16::mNG::3xFlag::AID]; daf-2(e1370); Ex[inx-1p::TIR1::mTur2::rps-27::NeoR +unc-122::GFP]*

OH16106 *sIs12199 [sra-25::GFP]; ot853 [daf-16::mNG::AID]; daf-2(e1370); otIs730 [Punc-11prom8+ehs-1prom7+rgef1prom2::TIR1::mTur2]*

OH16119 *ot853[daf-16::mNG::3xFlag::AID]; daf-2(e1370); otIs730[UPN::TIR1::mTur2]; otIs732[Psri-9::GFP]*

OH16176 *daf-2(e1370)*; *ot853[daf-16::mNeonGreen::3xFlag::AID]*; *otIs586*; *otEx7434[ceh-36_delASE::TIR1::mTur2]*

OH16496 *daf-7(e1372)*; *ot877[daf-3::TagRFP::3xFlag::AID]*; *Ex[Pift-20::TIR1::mRuby + Punc-122::GFP]*

OH16571 *otEx7585[Pges-1::daf-16(4A)::GFP; unc-122p::mCherry]*

OH16572 *otEx7586[Pges-1:: daf-16(4A)::GFP; unc-122p::mCherry]*

OH16573 *otEx7587[Pges-1:: daf-16(4A)::GFP; unc-122p::mCherry]*

OH16440 *ot853[daf-16::mNG::3xFlag::AID]; otEx7541[UPN::daf-2^DN^::EBFP2; unc-122p::mCherry]*

OH16441 *ot853[daf-16::mNG::3xFlag::AID]; otEx7542[UPN:: daf-2^DN^:: EBFP2; unc-122p::mCherry]*

OH16447 *ot853[daf-16::mNG::3xFlag::AID]; otEx7546[Pges-1:: daf-2^DN^:: EBFP2; unc-122p::mCherry]*

OH16449 *ot853[daf-16::mNG::3xFlag::AID]; otEx7548[Pges-1:: daf-2^DN^:: EBFP2; unc-122p::mCherry]*

OH16450 *ot853[daf-16::mNG::3xFlag::AID]; otEx7549[Pmyo-2::daf-2 daf-2^DN^:: EBFP2; unc-122p::mCherry]*

OH16451 *ot853[daf-16::mNG::3xFlag::AID; otEx7550[Pmyo-2:: daf-2^DN^:: EBFP2; unc-122p::mCherry]*

OH16665 *ot853[daf-16::mNG::3xFlag::AID]; daf-2(e1370);*

*otEx7628[ehs-1prom4::TIR1::mTur2, unc-122p::mCherry]*

OH16666 *ot853[daf-16::mNG::3xFlag::AID]; daf-2(e1370);*

*otEx7629[ehs-1prom4::TIR1::mTur2, unc-122p::mCherry]*

OH16752 *ot853[daf-16::mNG::3xFlag::AID]); daf-2(e1370);*

*otEx7668[unc-17prom4::TIR1::mTur2, unc-122p::mCherry]*

OH16754 *ot853[daf-16::mNG::3xFlag::AID]); daf-2(e1370);*

*otEx7669[unc-17prom4::TIR1::mTur2, unc-122p::mCherry]*

OH16766 *ot853[daf-16::mNG::3xFlag::AID]); daf-2(e1370);*

*otEx7670[eat-4prom7::TIR1::mTur2, unc-122p::mCherry]*

OH16767 *ot853[daf-16::mNG::3xFlag::AID]); daf-2(e1370);*

*otEx7671[eat-4prom7::TIR1::mTur2, unc-122p::mCherry]*

OH16694 *ot853[daf-16::mNG::AID]; daf-2(e1370); otTi1 [Pmyo-3::TIR1::mRuby];*

*trIs30[him-4p::MB::YFP + hmr-1b::DsRed2 + unc-129nsp::DsRed2]*

OH16896 *ot975[daf-16::mNeptune2.5::3xFlag::AID]; daf-2(e1370);*

*otIs730[UPN::TIR1::mTur2]; inx-2(ot906 [inx-2::sl2::yfp::h2b])*

OH16897 *ot975[daf-16::mNeptune2.5::3xFlag::AID]; daf-2(e1370);*

*otIs730[UPN::TIR1::mTur2]; otEx7112 [che-7_fosmid::sl2::yfp::h2b; pha-1(+); myo-2::bfp]*

OH16898 *ot853 [daf-16::mNG::3xFlag::AID]; daf-2(e1370); otEx7686[unc-25p4::TIR1::mTur2; unc-122::mCherry]*

OH16899 *ot853 [daf-16::mNG::3xFlag::AID]; daf-2(e1370); otEx7687[cho-1p14::TIR1::mTur2; unc-122::mCherry]*

### Generation of tissue-specific TIR1 lines

TIR1 cassette was amplified from pLZ31[pCFJ350modified-*Peft3*-TIR1-linker-mRuby- *unc-54* 3’UTR] (Zhang et al., 2015) and inserted into the miniMos plasmid pCFJ910 (Frokjaer-Jensen et al., 2014) carrying the neomycin resistance gene, NeoR, as a selectable marker. Tissue- or cell-specific promoters [UPN, *Pceh-36prom2*(ASE-), *Pmyo-3*, *Pdpy-7, Pinx-1*, *ehs--1prom4*, *eat-4prom7*, *unc-17prom4, unc-25prom4 and cho-1prom14*] were inserted in the resulting miniMos plasmid using restriction--free (RF) cloning or restriction/ligation. AWC-specific promoter *ceh-36prom2* is an 1852 bp (–1883 to –32) fragment of the *ceh-36* promoter with a mutated ASE motif (Etchberger et al., 2009). The UPN promoter was developed by Eviatar Yemini (Yemini et al., 2019) by concatenating three promoter fragments of panneuronally expressed genes: *unc-11prom8*, *ehs-1prom7* and *rgef-1prom2* (Stefanakis et al., 2015). *ehs-1prom4* is a 62 bp (-332 to -271) fragment of the *ehs-1* promoter that is expressed in all pharyngeal neurons (weak expression in I1 and I4)(Stefanakis et al., 2015). *eat--4prom7* is a 587 bp (-5,038 to -4,452) fragment of the *eat-4* promoter that is expressed in glutamatergic pharyngeal neurons I5 and M3 (weak expression in M5 and MI)(Serrano-Saiz et al., 2020). *unc-17prom4* is an 837 bp (-822 to +15) fragment of the *unc-17* promoter that is expressed in cholinergic pharyngeal neurons I1, M1, M2, M4, M5 and MC (weak expression in I3; also expressed in AIY, IL2, RIH and RIR)(Serrano-Saiz et al., 2020). *unc-25prom4* is a 188bp (-188 to 0) fragment of the *unc-25* cis-regulatory region that is expressed in GABAergic D-type motor neurons (Serrano-Saiz et al., 2020). *cho-1prom14* is a 400bp (-3700 to - 3300) fragment of the *cho-1* cis-regulatory region that is expressed in cholinergic RMDD, RMDV, SMD and SIA motor neurons (Serrano-Saiz et al., 2020).

For the *UPN::TIR1*, *AWC::TIR1*, *Pdpy-7::TIR1*, *ehs-1prom4*, *eat-4prom7* and *unc-17prom4* constructs, the fluorophore tagging TIR1 in the original plasmid (pLZ31) was replaced with mTurquoise2 that was amplified from pDD315 (mTurquoise2^SEC^2xHA), a gift from Daniel Dickinson (Addgene plasmid #73343 http://n2t.net/addgene:73343; RRID:Addgene_73343).

It should be noted that while single-copy TIR1 lines for non-neuronal tissues have resulted in specific protein depletion and quantifiable and specific phenotypes, we found that single-copy neuronal drivers of TIR1 were insufficient in that regard. Therefore, we used integrated or extrachromosomal multicopy arrays of TIR1 for neuronal expression (panneuronal or neuron type-specific, i.e., AIB-, AWC- and pharyngeal neuron-specific).

### Conditional alleles of *daf-16, daf-3* and *daf-12*

AID was amplified form pLZ29[pCFJ350modified-*Peft3*-AID(71-114)EmGFP-*unc-54-*3’UTR] (Zhang et al., 2015) and inserted into pDD268 and pDD284 (Dickinson et al., 2015). These plasmids were used for SEC-mediated CRISPR insertion of FP::AID tags into genomic loci of *daf-16*, *daf-12* and *daf-3* (Dickinson et al., 2015).

### Generation of *daf-2^DN^* and *daf-16(4A)* constructs

To create *daf-2^DN^*, a dominant negative version of the insulin/IGF-like receptor *daf-2*, the cDNA for the *daf-2 a* isoform was modified using RF cloning to replace the sequence encoding the tyrosine kinase and C-terminus domains (corresponding to amino acids 1,246 to 1,846) with the blue fluorescent protein *ebfp2* sequence. This *daf-2^DN^* sequence was subsequently cloned downstream of tissue-specific promoters (*Pmyo-2*, UPN, *Pges--1*) and upstream of *unc-54* 3’UTR using RF cloning.

For creating an insulin-signaling-independent form of *daf-16*, the cDNA for the *c* isoform of *daf-16* (referred to as the *daf-16a* in (Lee et al., 2001)) was synthesized *de novo* and four Ser/Thr residues were replaced with alanine (T54A, S240A, T242A, S314A). This *daf-16(4A)* sequence was subsequently translationally fused upstream of the *gfp* sequence of pPD95.75 plasmid and the entire construct was cloned downstream of the intestine-specific promoter *Pges-1* using RF cloning.

### Auxin treatment

Auxin treatment was performed by transferring worms to bacteria-seeded plates containing auxin. The natural auxin indole-3-acetic acid (IAA) was purchased from Alfa Aesar (#A10556). A 400 mM stock solution in ethanol was prepared and stored at 4°C for up to one month. NGM (Nematode Growth Medium) agar plates with fully grown OP50 bacterial lawn were coated with the auxin stock solution to a final concentration of 4 mM and used on the same day. All plates in the experiments were protected from light because of the light-sensitivity of auxin.

### Oil Red O staining

Intestinal lipid staining with Oil Red O was performed according to a protocol by (He, 2012).

### Lifespan analysis

Lifespan assays for strains with tissue-specific DAF-16/FoxO depletion were performed using the multi-well WorMotel platform (Churgin et al., 2017). We prepared 2 to 4 independent replicates of each experiment, with total N=40 to 80 worms per condition. More specifically, the following replicates were tested for each genotype: 1) *daf-2(e1370)*: 3 independent experiments, N=20 per condition; 2) *daf-2(e1370); daf-1::AID*: 4 independent experiments, n=20 per condition; 3) *daf-2(e1370); daf-16::AID; Peft-3::TIR1*: 4 independent experiments, n=20 per condition; 4) *daf-2(e1370); daf-1::AID; Pmyo-2:TIR1*: 2 independent experiments, n=20 per condition; 5) *daf-2(e1370); daf-16::AID; Pmyo-3:TIR1:* 2 independent experiments, n=20 per condition; 6) *daf-2(e1370); daf-16::AID; Pges-1:TIR1*: 2 independent experiments, n=20 per condition.

Images of worms in the WorMotel were acquired using static imaging systems (Churgin and Fang-Yen, 2015) with illumination from a blue LED array applied twice per day to stimulate movement. Survival curves were calculated as previously described (Churgin et al., 2017).

### Microscopy

Worms were anesthetized using 100 mM sodium azide (NaN_3_) and mounted on 5% agarose pads on glass slides. Images were acquired as Z-stacks of ∼0.7 μm-thick slices with Zen software (ZEN Digital Imaging for Light Microscopy, RRID:SCR_013672) using a Zeiss LSM700 confocal microscope. Images were reconstructed via maximum intensity Z-projection of 2-10 μm Z-stacks using Zen software.

### Automated worm tracking experiments

Automated single worm tracking was performed using WormTracker 2.0 system (Yemini et al., 2013) at 25°C. Animals were recorded for 5 min to ensure sufficient sampling of locomotion-related behavioral features. Dauer and non-dauer animals were placed on uncoated NGM plates that were kept at 25°C before recording. To minimize any potential bias arising due to dauer stage-specific prolonged bouts of spontaneous pausing (Gaglia and Kenyon, 2009), we only selected animals that were actively moving at the beginning of the assay. To avoid potential variability arising due to fluctuations in room conditions, all strains that were compared in a single experiment were recorded on the same day.

### Pharyngeal pumping rate assays

Constitutive dauer animals on an NGM plate containing OP50 bacteria were observed under 50x objective lens of a Nikon Eclipse E400 upright microscope equipped with DIC optics. The number of pharyngeal pumps in a 2 min period was measured for at least 15 animals per condition. For the N2 and *daf-16(m26)* strains that were not dauer-constitutive, dauer animals were isolated from a starved plate after treatment with 1% SDS for 30 min. SDS-resistant dauer individuals were picked and transferred to a fresh NGM plate seeded with a uniform thin layer of OP50 bacteria and allowed to acclimate for 15 min. Subsequently, pharyngeal pumps were scored (as described above) within 40 min of placing the dauer animals on the OP50 lawn plate.

For measurement of pharyngeal pumping rate in adults, day 1 adult worms grown at 25°C from hatching were picked and transferred from an uncrowded non-starved plate to a fresh NGM plate seeded with a uniform thin layer of OP50 bacteria. Worms were allowed to recover for 10 min and were subsequently visualized under 20x objective lens of a Nikon Eclipse E400 upright microscope. The number of pharyngeal pumps in a 1 min period was measured for at least 15 animals per condition. All pharyngeal pumping assays were performed at room temperature.

## ACKNOWLEDGEMENTS

We thank Chi Chen for generating transgenic strains, Tulsi Patel for originally noting the *gcy-5* expression change in dauers, Piali Sengupta and Michael O’Donnell for comments on the manuscript. This work was supported by NIH R21NS115442 and the Howard Hughes Medical Institute.

## SUPPLEMENTARY FIGURES/TABLES

**Fig.S1.**
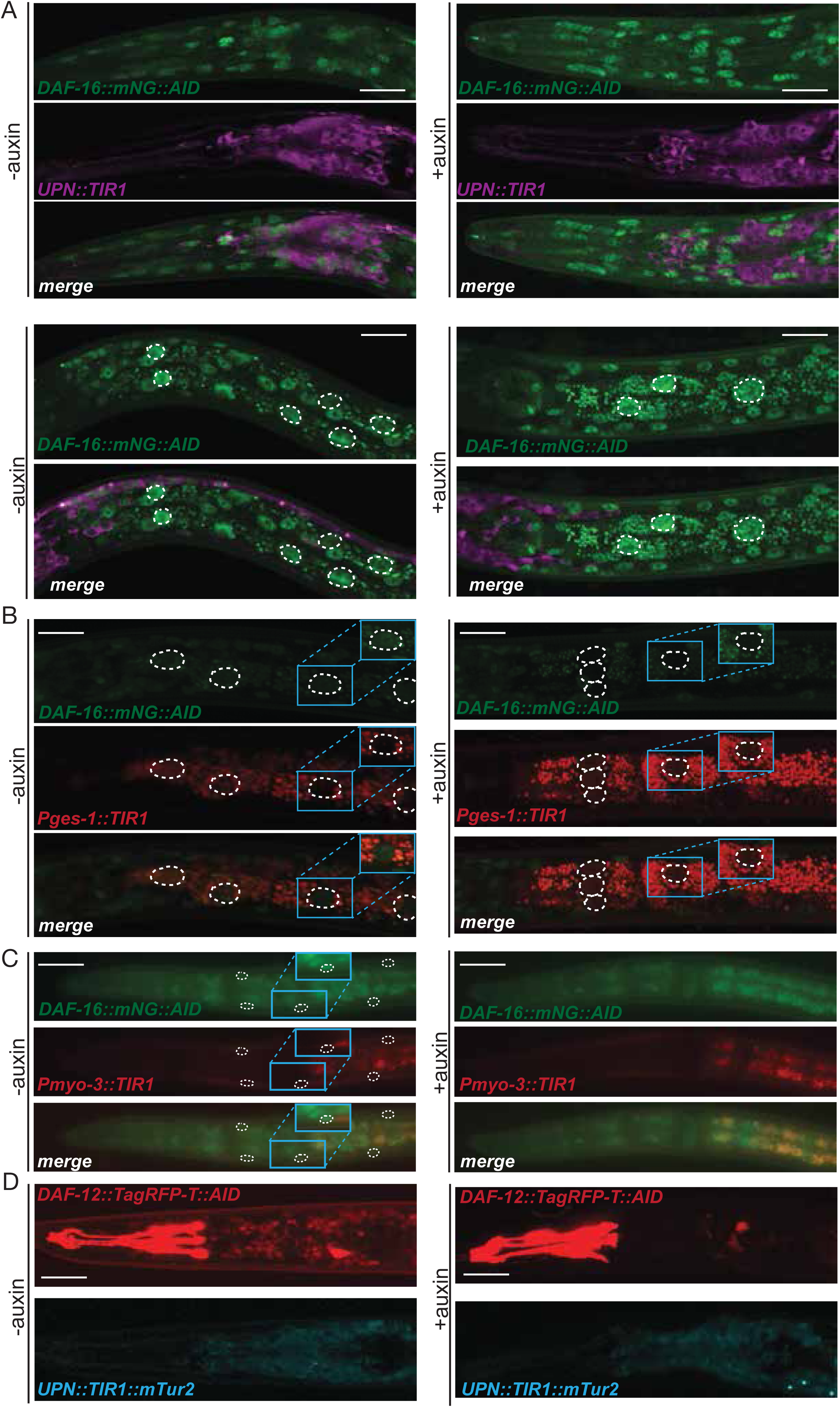
**GFP images indicating successful removal of TFs after auxin treatment** (A) Confocal images of dauers with panneuronal depletion of DAF-16/FoxO under control and auxin treatment conditions. Scale bar, 20 µm. (B) Confocal images of dauers with intestinal depletion of DAF-16/FoxO under control and auxin treatment conditions. Insets show an example of an intestinal nucleus, magnified and enhanced to boost fluorescence signal (both in control and auxin-treated conditions). Scale bar, 20 µm. (C) Epifluorescent images of dauers with body wall muscle depletion of DAF-16/FoxO under control and auxin treatment conditions. Insets show an example of a muscle nucleus, magnified and enhanced to boost fluorescence signal. Scale bar, 20 µm. (D) Confocal images of dauers with panneuronal depletion of DAF-12/VDR under control and auxin treatment conditions. Pharyngeal expression is from a co-injection marker *inx-6prom 18::TagRFP*. Scale bar, 20 µm.

**Fig.S2.**
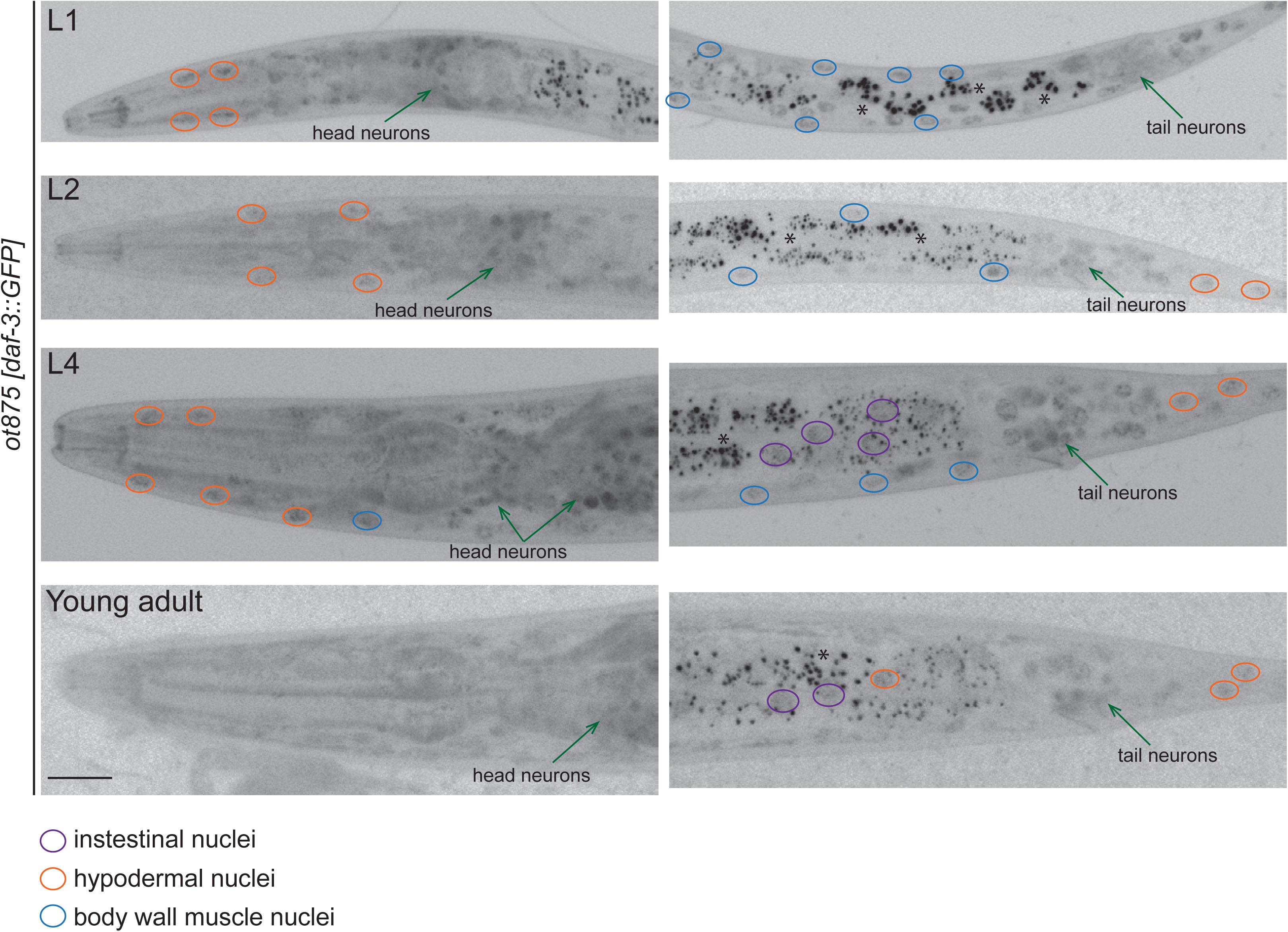
**DAF-3::GFP expression in starved larvae** Expression of the *daf-3::GFP* CRISPR allele at different stages in development in starved conditions. Anterior is to the left on all images. Scale bar, 20 µm (same for all images).

**Fig.S3.**
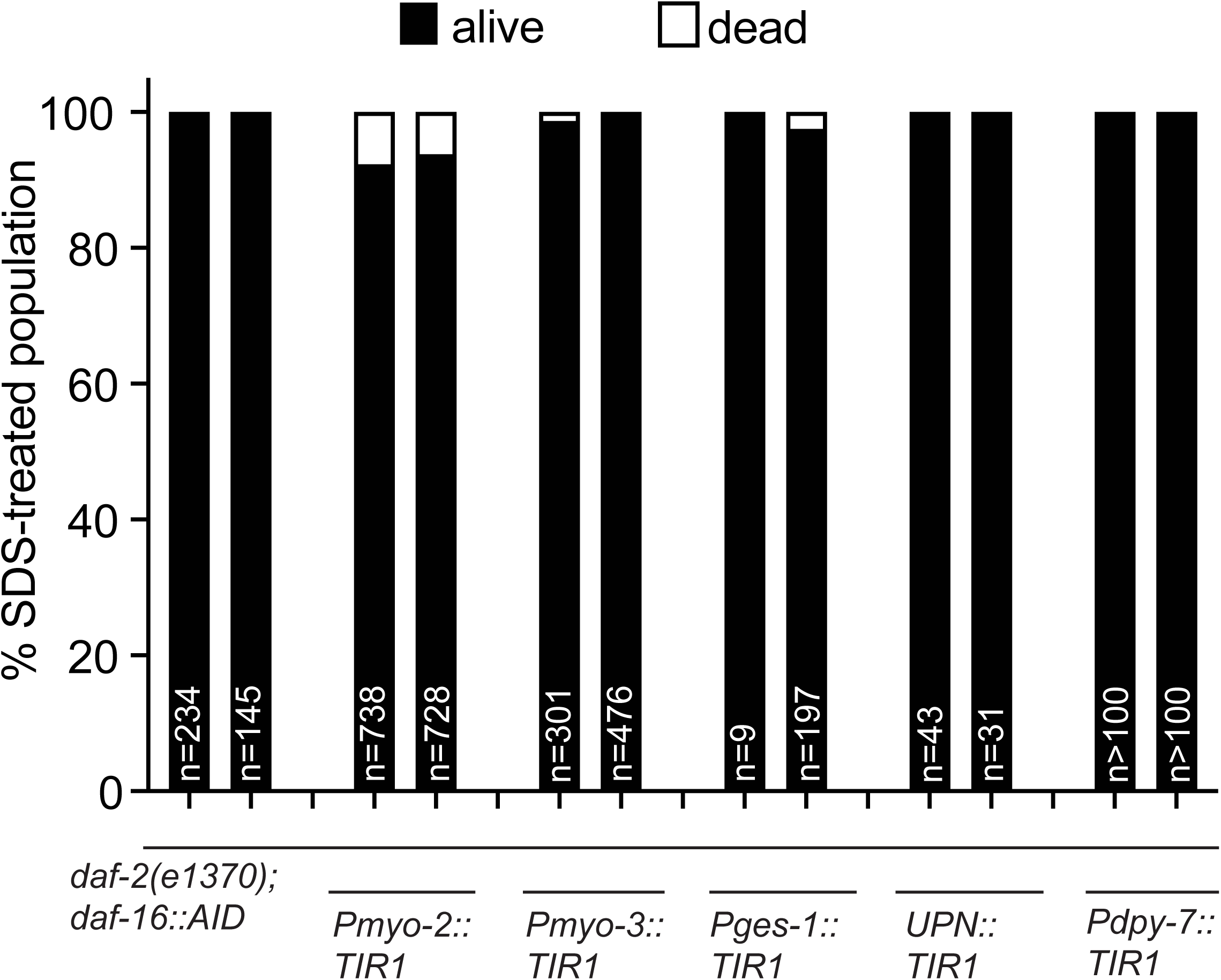
**Quantification of SDS-resistance of dauers of the conditional strains.** Worms were washed off NGM plates (control and auxin-treated) 3 days after rearing at 25°C and incubated in 1% (m/v) solution of SDS for 30 min, with continuous gentle shaking. After washing with water and M9 buffer, the worms were plated on fresh plates and scored as alive if moving.

**Fig.S4:**
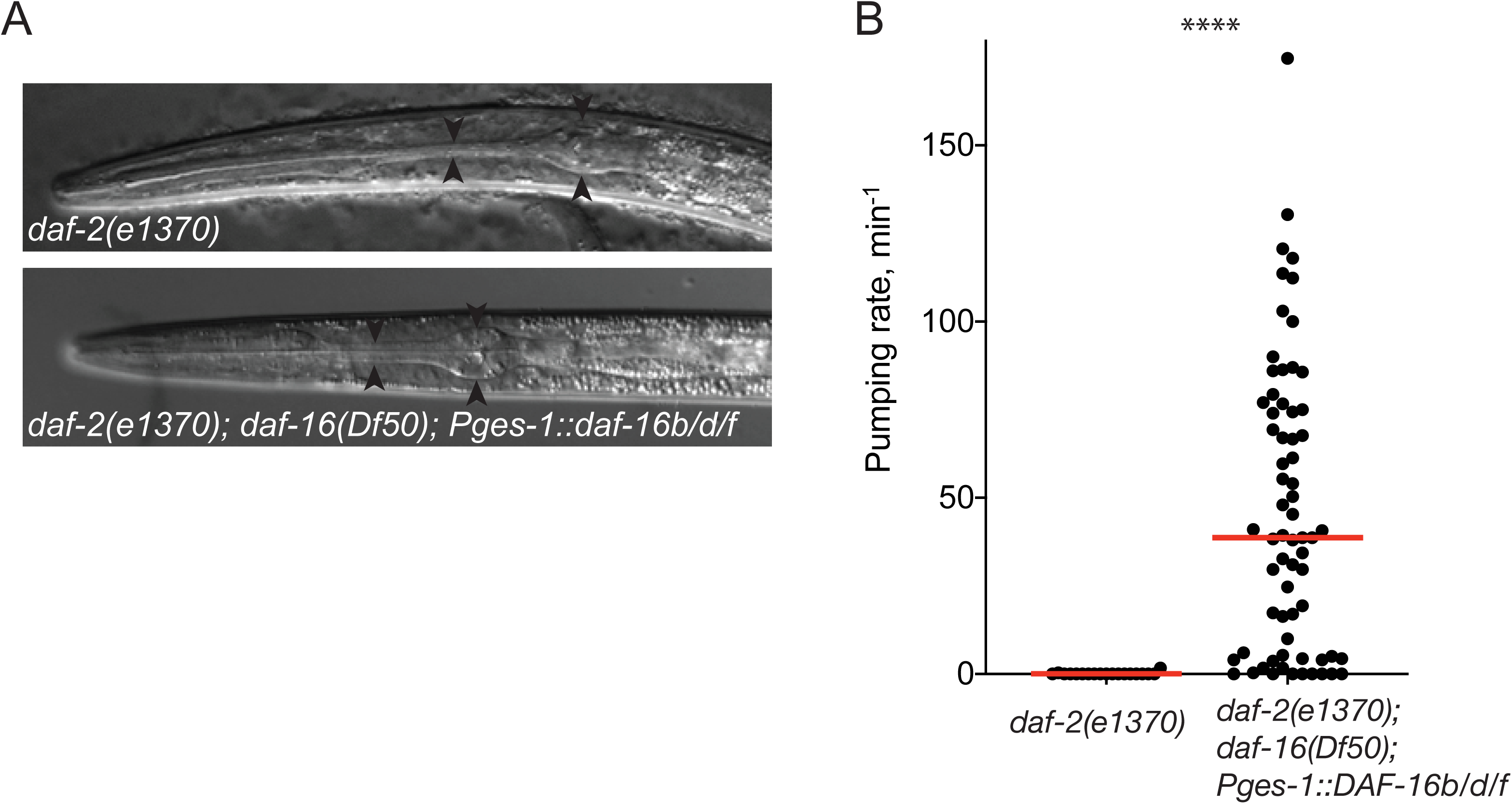
**Intestinal rescue of daf-16 results in incomplete rescue of the Daf-c phenotype of *daf-2*; *daf-16* mutants** (A) DIC images of a *daf-2(e1370)* dauer and a dauer-like larva with intestinal rescue of *daf-16b/d/f* isoforms in the *daf-2(e1370); daf-16(Df50)* background. Note the deviation from the wildtype filariform pharynx morphology in the latter case. (B) Quantification of pharyngeal activity of the strains depicted in (**A**).

**Fig.S5:**
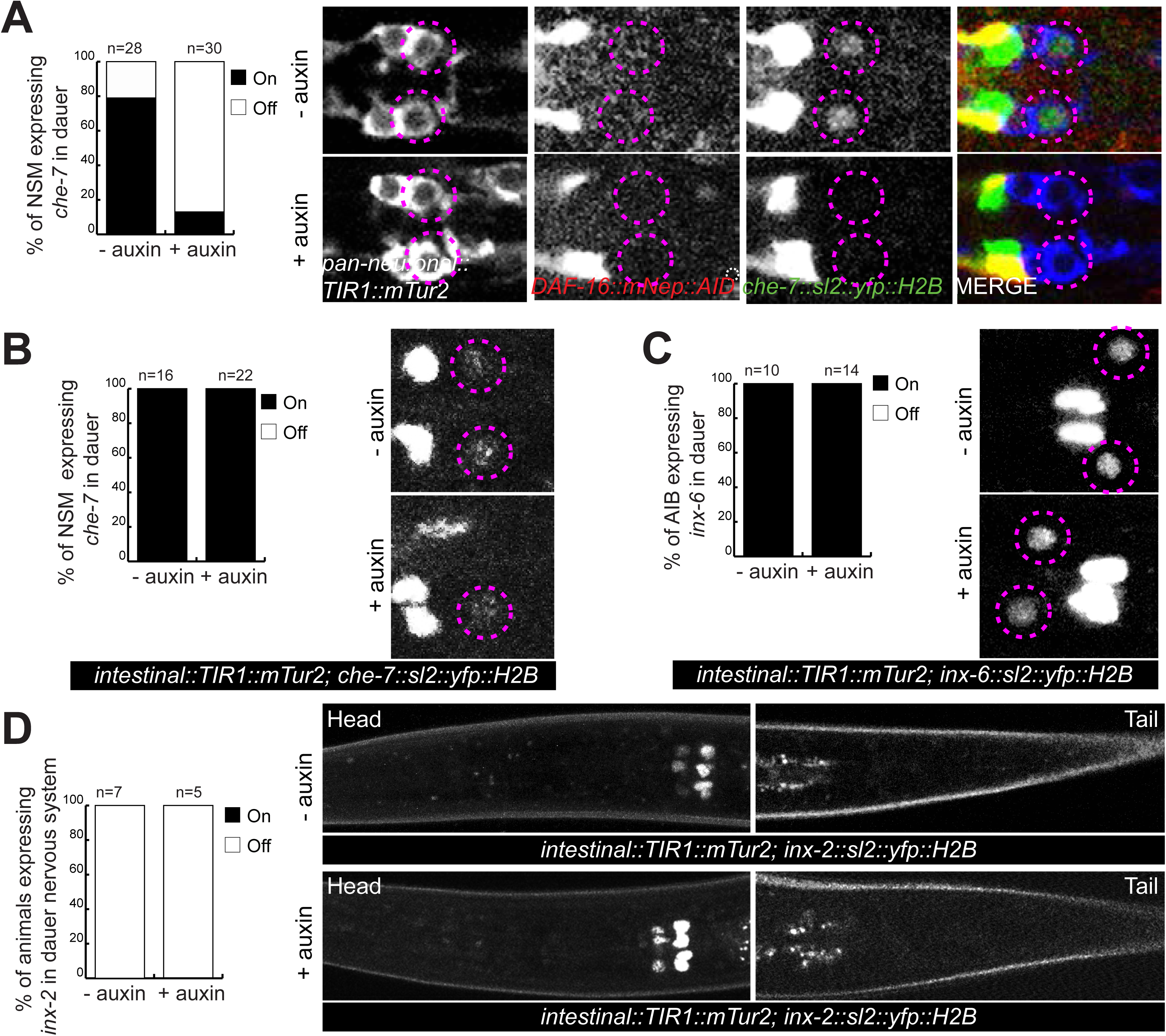
**Dauer-specific expression change of electrical synapse components is independent of intestinal DAF-16/FoxO activity.** (A) A *che-7* reporter (*otEx7112)* expression is gained in NSM neurons in dauer. This dauer-specific *che-7* expression is lost upon panneuronal depletion of DAF-16/FoxO in auxin-treated dauers. (B) Dauer-specific *che-7* expression in NSM is unaffected upon intestinal depletion of DAF-16/FoxO in auxin-treated dauers. (C) Expression of an *inx-6* reporter allele *(ot804)* is gained in AIB neurons in dauer. This dauer-specific *inx-6* expression in AIB neurons is unaffected upon intestinal depletion of daf-16/FoxO in auxin-treated dauers. (D) Expression of an *inx-2* reporter allele (*ot906)* is downregulated in multiple neurons in dauer. This dauer-specific downregulation of *inx-6* expression is unaffected upon intestinal depletion of DAF-16/FoxO in auxin-treated dauers.

**Fig.S6:**
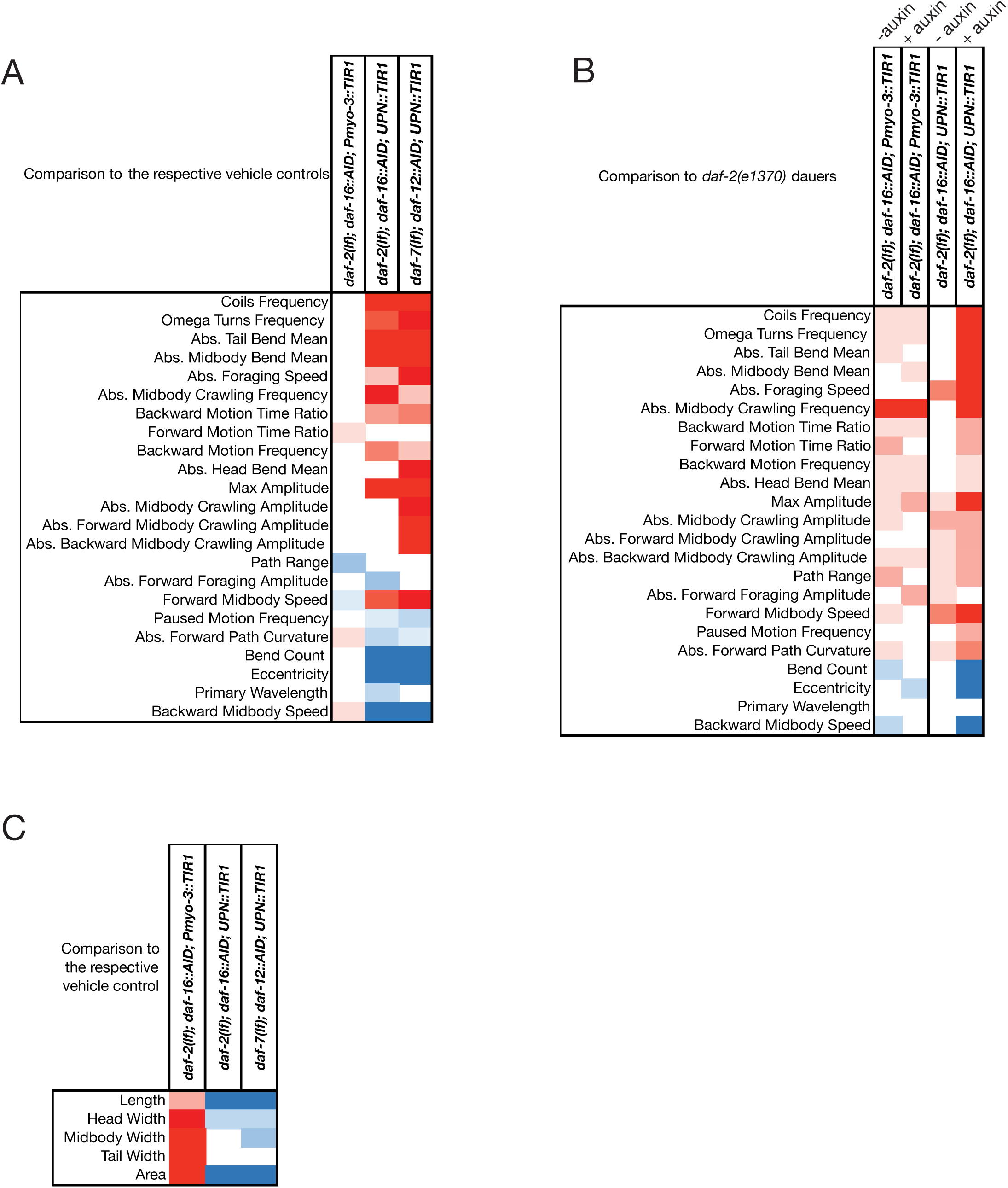
**(A)** Locomotory features of dauers with DAF-16 depleted from neurons and body wall muscle and DAF-12 depleted from neurons, as compared to their respective vehicle controls. (**B)** Locomotory features of dauers with DAF-16 depleted from neurons and body wall muscle, as compared to *daf-2(e1370)* control dauers. (**C**) Effect of DAF-16 depletion from body wall muscle on dauer morphology

**Fig.S7:**
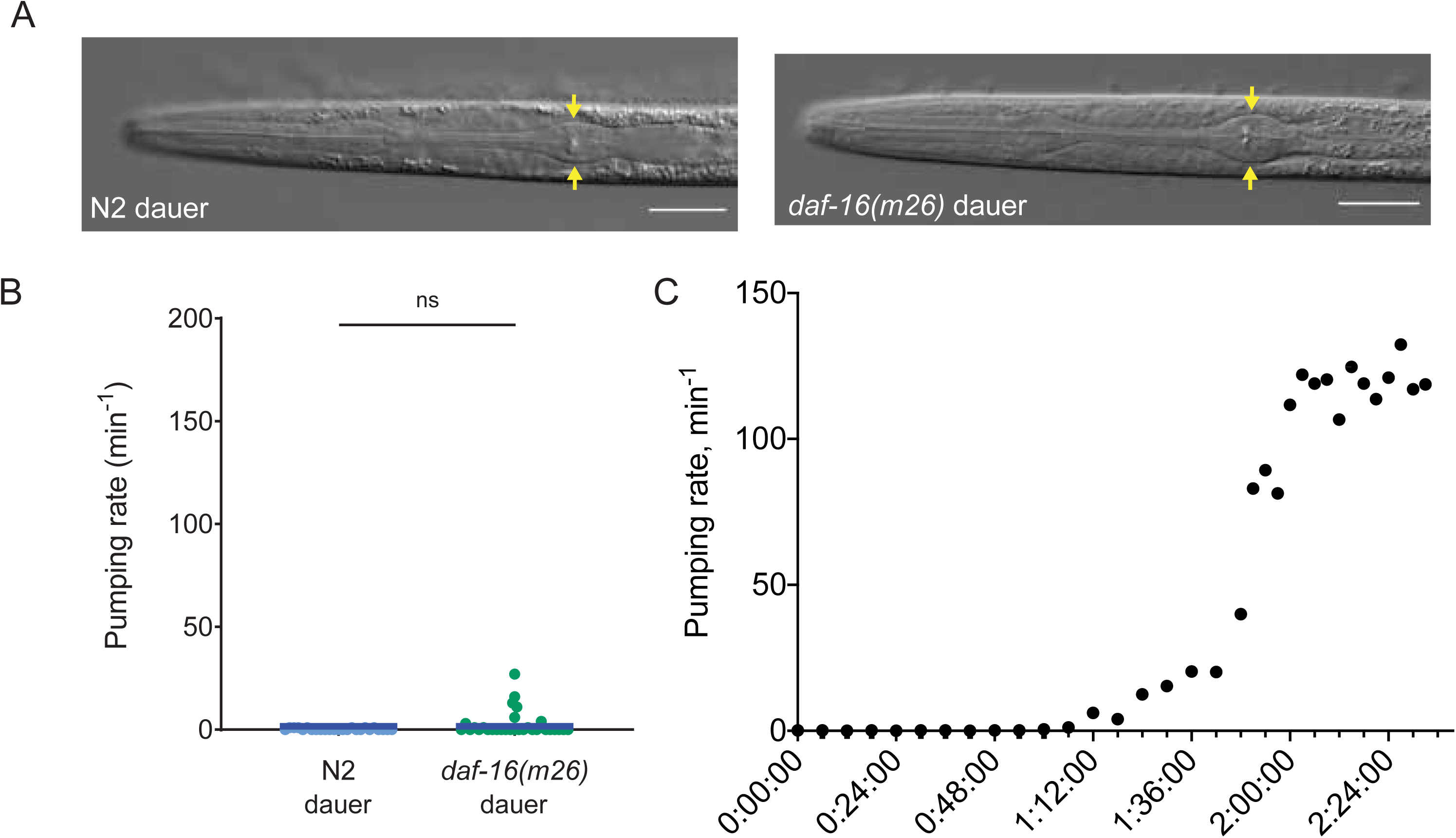
**Pharyngeal constriction and pharyngeal pumping can be decoupled** (A) Pharyngeal morphology in wild-type (N2) and *daf-16(m26)* starvation-induced dauers. Yellow arrows show width of the terminal bulb of the pharynx. Scale bars: 20 μm. (B) Pharyngeal pumping rate in N2 and *daf-16(m26)* starvation-induced dauer animals. Blue horizontal line represents median for ≥ 25 animals per genotype. ns indicates *p* = 0.36 in two-tailed Mann-Whitney test. (C) Pharyngeal pumping rate in wild-type animals recovering from starvation-induced dauer stage.

## Notes

### Competing Interest Statement

The authors have declared no competing interest.

## Bibliography

Aghayeva, U., Bhattacharya, A., and Hobert, O. (2020). A panel of fluorophore-tagged daf-16 alleles. MicroPubl Biol 2020.

Albert, P.S., and Riddle, D.L. (1983). Developmental alterations in sensory neuroanatomy of the Caenorhabditis elegans dauer larva. The Journal of comparative neurology 219, 461–481.

Albertson, D.G., and Thomson, J.N. (1976). The pharynx of Caenorhabditis elegans. Philos Trans R Soc Lond B Biol Sci 275, 299–325.

Androwski, R.J., Flatt, K.M., and Schroeder, N.E. (2017). Phenotypic plasticity and remodeling in the stress-induced Caenorhabditis elegans dauer. Wiley interdisciplinary reviews Developmental biology 6.

Antebi, A. (2013). Steroid regulation of C. elegans diapause, developmental timing, and longevity. Curr Top Dev Biol 105, 181–212.

Antebi, A., Culotti, J.G., and Hedgecock, E.M. (1998). daf-12 regulates developmental age and the dauer alternative in Caenorhabditis elegans. Development 125, 1191–1205.

Antebi, A., Yeh, W.H., Tait, D., Hedgecock, E.M., and Riddle, D.L. (2000). daf-12 encodes a nuclear receptor that regulates the dauer diapause and developmental age in C. elegans. Genes Dev 14, 1512–1527.

Apfeld, J., and Kenyon, C. (1998). Cell nonautonomy of C. elegans daf-2 function in the regulation of diapause and life span. Cell 95, 199–210.

Avery, L., and Thomas, J.H. (1997). Feeding and Defecation. In Celegans II, D.L. Riddle, T. Blumenthal, B.J. Meyer, and J.R. Priess, eds. (Cold Spring Harbor Laboratory Press), pp. 679–716.

Bansal, A., Kwon, E.S., Conte, D., Jr., Liu, H., Gilchrist, M.J., MacNeil, L.T., and Tissenbaum, H.A. (2014). Transcriptional regulation of Caenorhabditis elegans FOXO/DAF-16 modulates lifespan. Longev Healthspan 3, 5.

Bhattacharya, A., Aghayeva, U., Berghoff, E.G., and Hobert, O. (2019). Plasticity of the Electrical Connectome of C. elegans. Cell 176, 1174–1189 e1116.

Blau, H.M. (1992). Differentiation requires continuous active control. Annu Rev Biochem 61, 1213–1230.

Burnell, A.M., Houthoofd, K., O’Hanlon, K., and Vanfleteren, J.R. (2005). Alternate metabolism during the dauer stage of the nematode Caenorhabditis elegans. Exp Gerontol 40, 850–856.

Cassada, R.C., and Russell, R.L. (1975). The dauerlarva, a post-embryonic developmental variant of the nematode Caenorhabditis elegans. Dev Biol 46, 326–342.

Churgin, M.A., and Fang-Yen, C. (2015). An Imaging System for C. elegans Behavior. Methods Mol Biol 1327, 199–207.

Churgin, M.A., Jung, S.K., Yu, C.C., Chen, X., Raizen, D.M., and Fang-Yen, C. (2017). Longitudinal imaging of Caenorhabditis elegans in a microfabricated device reveals variation in behavioral decline during aging. eLife 6.

Cook, S.J., Crouse, C.M., Yemini, E., Hall, D.H., Emmons, S.W., and Hobert, O. (2020). The connectome of the Caenorhabditis elegans pharynx. The Journal of comparative neurology.

Cui, X., Gooch, H., Petty, A., McGrath, J.J., and Eyles, D. (2017). Vitamin D and the brain: Genomic and non-genomic actions. Mol Cell Endocrinol 453, 131–143.

Dickinson, D.J., Pani, A.M., Heppert, J.K., Higgins, C.D., and Goldstein, B. (2015). Streamlined Genome Engineering with a Self-Excising Drug Selection Cassette. Genetics 200, 1035–1049.

Dixon, S.J., Alexander, M., Chan, K.K., and Roy, P.J. (2008). Insulin-like signaling negatively regulates muscle arm extension through DAF-12 in Caenorhabditis elegans. Dev Biol 318, 153–161.

Dixon, S.J., and Roy, P.J. (2005). Muscle arm development in Caenorhabditis elegans. Development 132, 3079–3092.

Duret, L., Guex, N., Peitsch, M.C., and Bairoch, A. (1998). New insulin-like proteins with atypical disulfide bond pattern characterized in Caenorhabditis elegans by comparative sequence analysis and homology modeling. Genome Res 8, 348–353.

Etchberger, J.F., Flowers, E.B., Poole, R.J., Bashllari, E., and Hobert, O. (2009). Cis-regulatory mechanisms of left/right asymmetric neuron-subtype specification in C. elegans. Development 136, 147–160.

Fernandez, A.M., and Torres-Aleman, I. (2012). The many faces of insulin-like peptide signalling in the brain. Nat Rev Neurosci 13, 225–239.

Ferrario, C.R., and Reagan, L.P. (2018). Insulin-mediated synaptic plasticity in the CNS: Anatomical, functional and temporal contexts. Neuropharmacology 136, 182–191.

Fielenbach, N., and Antebi, A. (2008). C. elegans dauer formation and the molecular basis of plasticity. Genes Dev 22, 2149–2165.

Frokjaer-Jensen, C., Davis, M.W., Sarov, M., Taylor, J., Flibotte, S., LaBella, M., Pozniakovsky, A., Moerman, D.G., and Jorgensen, E.M. (2014). Random and targeted transgene insertion in Caenorhabditis elegans using a modified Mos1 transposon. Nat Methods 11, 529–534.

Gaglia, M.M., and Kenyon, C. (2009). Stimulation of movement in a quiescent, hibernation-like form of Caenorhabditis elegans by dopamine signaling. J Neurosci 29, 7302–7314.

Golden, J.W., and Riddle, D.L. (1982). A pheromone influences larval development in the nematode Caenorhabditis elegans. Science 218, 578–580.

Gottlieb, S., and Ruvkun, G. (1994). daf-2, daf-16 and daf-23: genetically interacting genes controlling Dauer formation in Caenorhabditis elegans. Genetics 137, 107–120.

Hall, S.E., Beverly, M., Russ, C., Nusbaum, C., and Sengupta, P. (2010). A cellular memory of developmental history generates phenotypic diversity in C. elegans. Curr Biol 20, 149–155.

He, F. (2012). Oil Red O Staining of Fixed Worm. Bio-101 e230.

Hobert, O. (2016). Terminal Selectors of Neuronal Identity. Curr Top Dev Biol 116, 455–475.

Hung, W.L., Wang, Y., Chitturi, J., and Zhen, M. (2014). A Caenorhabditis elegans developmental decision requires insulin signaling-mediated neuron-intestine communication. Development 141, 1767–1779.

Hunt-Newbury, R., Viveiros, R., Johnsen, R., Mah, A., Anastas, D., Fang, L., Halfnight, E., Lee, D., Lin, J., Lorch, A., et al. (2007). High-throughput in vivo analysis of gene expression in Caenorhabditis elegans. PLoS Biol 5, e237.

Inoue, T., and Thomas, J.H. (2000). Targets of TGF-beta signaling in Caenorhabditis elegans dauer formation. Dev Biol 217, 192–204.

Jeong, M.H., Kawasaki, I., and Shim, Y.H. (2010). A circulatory transcriptional regulation among daf-9, daf-12, and daf-16 mediates larval development upon cholesterol starvation in Caenorhabditis elegans. Dev Dyn 239, 1931–1940.

Keane, J., and Avery, L. (2003). Mechanosensory inputs influence Caenorhabditis elegans pharyngeal activity via ivermectin sensitivity genes. Genetics 164, 153–162.

Kenyon, C., Chang, J., Gensch, E., Rudner, A., and Tabtiang, R. (1993). A C. elegans mutant that lives twice as long as wild type. Nature 366, 461–464.

Kimura, K.D., Tissenbaum, H.A., Liu, Y., and Ruvkun, G. (1997). daf-2, an insulin receptor-like gene that regulates longevity and diapause in Caenorhabditis elegans [see comments]. Science 277, 942–946.

Lee, H., Choi, M.K., Lee, D., Kim, H.S., Hwang, H., Kim, H., Park, S., Paik, Y.K., and Lee, J. (2012). Nictation, a dispersal behavior of the nematode Caenorhabditis elegans, is regulated by IL2 neurons. Nat Neurosci 15, 107–112.

Lee, J.S., Shih, P.Y., Schaedel, O.N., Quintero-Cadena, P., Rogers, A.K., and Sternberg, P.W. (2017). FMRFamide-like peptides expand the behavioral repertoire of a densely connected nervous system. Proc Natl Acad Sci U S A 114, E10726–E10735.

Lee, R.Y., Hench, J., and Ruvkun, G. (2001). Regulation of C. elegans DAF-16 and its human ortholog FKHRL1 by the daf-2 insulin-like signaling pathway. Curr Biol 11, 1950–1957.

Lemieux, G.A., and Ashrafi, K. (2015). Neural Regulatory Pathways of Feeding and Fat in Caenorhabditis elegans. Annu Rev Genet 49, 413–438.

Lewis, J.A., Fleming, J.T., McLafferty, S., Murphy, H., and Wu, C. (1987). The levamisole receptor, a cholinergic receptor of the nematode Caenorhabditis elegans. Mol Pharmacol 31, 185–193.

Libina, N., Berman, J.R., and Kenyon, C. (2003). Tissue-specific activities of C. elegans DAF-16 in the regulation of lifespan. Cell 115, 489–502.

Lin, K., Dorman, J.B., Rodan, A., and Kenyon, C. (1997). daf-16: An HNF-3/forkhead family member that can function to double the life-span of Caenorhabditis elegans. Science 278, 1319–1322.

Lin, K., Hsin, H., Libina, N., and Kenyon, C. (2001). Regulation of the Caenorhabditis elegans longevity protein DAF-16 by insulin/IGF-1 and germline signaling. Nat Genet 28, 139–145.

Lindblom, T.H., Pierce, G.J., and Sluder, A.E. (2001). A C. elegans orphan nuclear receptor contributes to xenobiotic resistance. Curr Biol 11, 864–868.

Matyash, V., Entchev, E.V., Mende, F., Wilsch-Brauninger, M., Thiele, C., Schmidt, A.W., Knolker, H.J., Ward, S., and Kurzchalia, T.V. (2004). Sterol-derived hormone(s) controls entry into diapause in Caenorhabditis elegans by consecutive activation of DAF-12 and DAF-16. PLoS Biol 2, e280.

McEwen, B.S. (2010). Stress, sex, and neural adaptation to a changing environment: mechanisms of neuronal remodeling. Ann N Y Acad Sci 1204 Suppl, E38–59.

Melendez, A., Talloczy, Z., Seaman, M., Eskelinen, E.L., Hall, D.H., and Levine, B. (2003). Autophagy genes are essential for dauer development and life-span extension in C. elegans. Science 301, 1387–1391.

Mills, J.C., Stanger, B.Z., and Sander, M. (2019). Nomenclature for cellular plasticity: are the terms as plastic as the cells themselves? EMBO J 38, e103148.

Motola, D.L., Cummins, C.L., Rottiers, V., Sharma, K.K., Li, T., Li, Y., Suino-Powell, K., Xu, H.E., Auchus, R.J., Antebi, A., et al. (2006). Identification of ligands for DAF-12 that govern dauer formation and reproduction in C. elegans. Cell 124, 1209–1223.

Murphy, C.T., McCarroll, S.A., Bargmann, C.I., Fraser, A., Kamath, R.S., Ahringer, J., Li, H., and Kenyon, C. (2003). Genes that act downstream of DAF-16 to influence the lifespan of Caenorhabditis elegans. Nature 424, 277–283.

Nolan, K.M., Sarafi-Reinach, T.R., Horne, J.G., Saffer, A.M., and Sengupta, P. (2002). The DAF-7 TGF-beta signaling pathway regulates chemosensory receptor gene expression in C. elegans. Genes Dev 16, 3061–3073.

Ogg, S., Paradis, S., Gottlieb, S., Patterson, G.I., Lee, L., Tissenbaum, H.A., and Ruvkun, G. (1997). The Fork head transcription factor DAF-16 transduces insulin-like metabolic and longevity signals in C. elegans. Nature 389, 994–999.

Oh, S.W., Mukhopadhyay, A., Dixit, B.L., Raha, T., Green, M.R., and Tissenbaum, H.A. (2006). Identification of direct DAF-16 targets controlling longevity, metabolism and diapause by chromatin immunoprecipitation. Nat Genet 38, 251–257.

Ortiz, C.O., Faumont, S., Takayama, J., Ahmed, H.K., Goldsmith, A.D., Pocock, R., McCormick, K.E., Kunimoto, H., Iino, Y., Lockery, S., et al. (2009). Lateralized gustatory behavior of C. elegans is controlled by specific receptor-type guanylyl cyclases. Curr Biol 19, 996–1004.

Paradis, S., and Ruvkun, G. (1998). Caenorhabditis elegans Akt/PKB transduces insulin receptor-like signals from AGE-1 PI3 kinase to the DAF-16 transcription factor. Genes Dev 12, 2488–2498.

Patel, T., and Hobert, O. (2017). Coordinated control of terminal differentiation and restriction of cellular plasticity. eLife 6.

Patterson, G.I., Koweek, A., Wong, A., Liu, Y., and Ruvkun, G. (1997). The DAF-3 Smad protein antagonizes TGF-beta-related receptor signaling in the Caenorhabditis elegans dauer pathway. Genes Dev 11, 2679–2690.

Peckol, E.L., Troemel, E.R., and Bargmann, C.I. (2001). Sensory experience and sensory activity regulate chemosensory receptor gene expression in Caenorhabditis elegans. Proc Natl Acad Sci U S A 98, 11032–11038.

Procko, C., Lu, Y., and Shaham, S. (2011). Glia delimit shape changes of sensory neuron receptive endings in C. elegans. Development 138, 1371–1381.

Ren, P., Lim, C.S., Johnsen, R., Albert, P.S., Pilgrim, D., and Riddle, D.L. (1996). Control of C. elegans larval development by neuronal expression of a TGF-beta homolog. Science 274, 1389–1391.

Ritter, A.D., Shen, Y., Fuxman Bass, J., Jeyaraj, S., Deplancke, B., Mukhopadhyay, A., Xu, J., Driscoll, M., Tissenbaum, H.A., and Walhout, A.J. (2013). Complex expression dynamics and robustness in C. elegans insulin networks. Genome Res 23, 954–965.

Rottiers, V., Motola, D.L., Gerisch, B., Cummins, C.L., Nishiwaki, K., Mangelsdorf, D.J., and Antebi, A. (2006). Hormonal control of C. elegans dauer formation and life span by a Rieske-like oxygenase. Dev Cell 10, 473–482.

Schroeder, N.E., Androwski, R.J., Rashid, A., Lee, H., Lee, J., and Barr, M.M. (2013). Dauer-specific dendrite arborization in C. elegans is regulated by KPC-1/Furin. Curr Biol 23, 1527–1535.

Serrano-Saiz, E., Gulez, B., Pereira, L., Gendrel, M., Kerk, S.Y., Vidal, B., Feng, W., Wang, C., Kratsios, P., Rand, J.B., et al. (2020). Modular Organization of Cis-regulatory Control Information of Neurotransmitter Pathway Genes in Caenorhabditis elegans. Genetics 215, 665–681.

Sims, J.R., Ow, M.C., Nishiguchi, M.A., Kim, K., Sengupta, P., and Hall, S.E. (2016). Developmental programming modulates olfactory behavior in C. elegans via endogenous RNAi pathways. eLife 5.

Srinivasan, S. (2020). Neuroendocrine control of lipid metabolism: lessons from C. elegans. J Neurogenet, 1–7.

Stefanakis, N., Carrera, I., and Hobert, O. (2015). Regulatory Logic of Pan-Neuronal Gene Expression in C. elegans. Neuron 87, 733–750.

Thomas, J.H., Birnby, D.A., and Vowels, J.J. (1993). Evidence for parallel processing of sensory information controlling dauer formation in Caenorhabditis elegans. Genetics 134, 1105–1117.

Timsit, Y.E., and Negishi, M. (2007). CAR and PXR: the xenobiotic-sensing receptors. Steroids 72, 231–246.

Vidal, B., Aghayeva, U., Sun, H., Wang, C., Glenwinkel, L., Bayer, E.A., and Hobert, O. (2018). An atlas of Caenorhabditis elegans chemoreceptor expression. PLoS Biol 16, e2004218.

Vowels, J.J., and Thomas, J.H. (1992). Genetic analysis of chemosensory control of dauer formation in Caenorhabditis elegans. Genetics 130, 105–123.

Wolkow, C.A., Kimura, K.D., Lee, M.S., and Ruvkun, G. (2000). Regulation of C. elegans life-span by insulinlike signaling in the nervous system. Science 290, 147–150.

Yemini, E., Jucikas, T., Grundy, L.J., Brown, A.E., and Schafer, W.R. (2013). A database of Caenorhabditis elegans behavioral phenotypes. Nat Methods 10, 877–879.

Yemini, E., Lin, A., Nejatbakhsh, A., Varol, E., Sun, R., Mena, G.E., Samuel, A.D.T., Paninski, L., Venkatachalam, V., and Hobert, O. (2019). NeuroPAL: A Neuronal Polychromatic Atlas of Landmarks for Whole-Brain Imaging in C. elegans. bioRxiv.

Yu, S., Avery, L., Baude, E., and Garbers, D.L. (1997). Guanylyl cyclase expression in specific sensory neurons: a new family of chemosensory receptors. Proc Natl Acad Sci U S A 94, 3384–3387.

Zaret, K.S., and Carroll, J.S. (2011). Pioneer transcription factors: establishing competence for gene expression. Genes Dev 25, 2227–2241.

Zhang, L., Ward, J.D., Cheng, Z., and Dernburg, A.F. (2015). The auxin-inducible degradation (AID) system enables versatile conditional protein depletion in C. elegans. Development 142, 4374–4384.

Zhang, P., Judy, M., Lee, S.J., and Kenyon, C. (2013). Direct and indirect gene regulation by a life-extending FOXO protein in C. elegans: roles for GATA factors and lipid gene regulators. Cell Metab 17, 85–100.

